# Engineered dual-gastruloid model for foregut-pharyngeal mesoderm co-development

**DOI:** 10.64898/2026.09.25.753740

**Authors:** Giada Mura, Michiel Tawdarous, Patrícia Duarte, Florian Mueller, Alexandre Mayran, Ingo Burtscher, Heiko Lickert, Shahragim Tajbakhsh

**Affiliations:** Stem Cells & Development Unit, Institut Pasteur, CNRS UMR 3738; Photonic BioImaging (UTechS PBI), Quantitative RNA imaging, Institut Pasteur, 25 rue du Dr. Roux, Paris F-75015, France; Institut Cochin, Université Paris Cité, CNRS UMR8104, INSERM U1016, Paris, France; Institute of Diabetes and Regeneration Research, Helmholtz Center Munich, 85764 Neuherberg, Germany; German Center for Diabetes Research (DZD), Neuherberg, Germany; School of Medicine, Technical University of Munich (TUM), Munich, Germany

**Keywords:** Craniofacial, gastruloid, embryo model, Foxa2, Tbx1, Shh, Hedgehog signalling

## Abstract

During mammalian gastrulation, progenitors of craniofacial and cardiopharyngeal muscles arise from anterior mesoderm adjacent to foregut endoderm, and they are controlled by gene regulatory networks and signalling pathways distinct from those of trunk mesoderm. While *in vitro* models capture aspects of posterior and heart development, a spatially organised interface between foregut-like endoderm and cranial-mesoderm-like tissue has not been described. Here we introduce a dual-aggregate gastruloid model, formed by fusing two mouse embryonic stem cell aggregates patterned under distinct Nodal, FGF, and Wnt signalling. These structures reproducibly establish an anterior-posterior axis and position a Foxa2⁺/Sox17⁺/Cdh1⁺ foregut-like epithelium proximal to Pax1⁺/Pax9⁺ pharyngeal-like endoderm and Tbx1⁺ cranial-mesoderm-like cells, resembling the E8.5 pharyngeal region. Reciprocal chimeras and *Foxa2* loss-of-function indicate that Foxa2⁺ endoderm is required for *Tbx1* expression in adjacent anterior mesoderm. Pharmacological perturbation of Hedgehog disrupts *Foxa2* and *Tbx1* expression. The model provides a tractable platform for dissecting endoderm-mesoderm signalling and congenital craniofacial disease mechanisms.

## INTRODUCTION

Cell fate specification during gastrulation emerges through dynamic interactions between intrinsic transcriptional competence and extrinsic morphogen gradients. In mammals, lineage diversification along the primitive streak is regulated by spatiotemporal Nodal/Wnt gradients, where high Nodal activity promotes definitive endoderm formation (Vincent et al., 2003; Dunn et al., 2004), whereas combined Wnt and moderate Nodal signalling generates anterior mesoderm (Dunn et al., 2004; Probst et al., 2021; Robertson, 2014; Vincent et al., 2003). This anterior domain gives rise to craniofacial and cardiopharyngeal mesoderm (Kinder et al., 1999; Parameswaran and Tam, 1995; Probst et al., 2021; Tam et al., 1997), transient progenitors that will populate the face, neck, and heart, through progressive lineage restrictions as cells encounter new signalling neighbours, notably endoderm and neural crest cells (Lescroart et al., 2022; Tam et al., 1997; Tzahor and Evans, 2011). Trunk paraxial mesoderm and other posterior tissues emerge after the streak has extended and its signalling landscape has remodelled (Duarte et al., 2023; Nandkishore et al., 2018; Shih et al., 2008).

The earliest neighboring tissue interaction between cranial mesoderm and foregut endoderm is transient and difficult to access *in vivo*, yet these interactions are critical for pharyngeal arch development. Signals such as Sonic Hedgehog (Shh), FGF, and BMP emanate from the foregut endoderm to pattern adjacent mesoderm, specifying domains marked by Tbx1, a transcription factor whose loss underlies DiGeorge syndrome (Garg et al., 2001; Guzzetta et al., 2020; Litingtung et al., 1998; Piotrowski and Nüsslein-Volhard, 2000; Yamagishi et al., 2003). Despite decades of embryological insight, how these tissues first meet and reciprocally instruct each other remains incompletely understood.

Gastruloids, aggregates of pluripotent stem cells (PSCs) that self-organise under transient Wnt activation, recapitulate symmetry breaking and posterior patterning with high fidelity (Beccari et al., 2018; Moris et al., 2020; Van Den Brink et al., 2014), but their anterior development is limited: cardiopharyngeal mesoderm and foregut endoderm are under-represented, whereas trunk identities predominate (Miao et al., 2023; Van Den Brink et al., 2020; Veenvliet et al., 2020; Yamanaka et al., 2023). Modified protocols extend anteriorisation and representation. Aggregates formed without exogenous Wnt activation and subsequently exposed to a Wnt inhibitor co-derive anterior neural progenitors and posterior tissues (Girgin et al., 2021), and cardiogenic conditions yield gastruloids expressing cardiopharyngeal markers (Argiro et al., 2024; Rossi et al., 2021). Chimeric gastrula and gastruloids built from cells of two genotypes can dissect gene function (Wehmeyer et al., 2022). Gastruloid approaches have not, however, generated a stable interface at which foregut-like endoderm and cranial-mesoderm-like tissue lie adjacent and can be perturbed independently.

We therefore assembled two aggregates with different signalling histories into an engineered dual-aggregate gastruloid model (Martinez Arias et al., 2024; Matthews et al., 2021). One (EPI) is biased toward anterior/early streak identity by Activin and FGF (Brons et al., 2007; Hayashi et al., 2011; Kojima et al., 2014); the other (CHI) acquires posterior/late streak identity following a Wnt agonist pulse (Dias et al., 2025; Turner et al., 2014). On fusion, they form a composite structure spanning anterior and posterior domains, hereafter EPICHI which reproducibly generates an anterior Foxa2⁺/Sox17⁺ foregut-like epithelium juxtaposed with Mesp1⁺ mesoderm and in which either compartment can be perturbed genetically or pharmacologically.

Here, we show that EPICHIs reproducibly juxtapose foregut-like endoderm and cranial-mesoderm-like tissue at the anterior pole, capturing a Hedgehog–Foxa2–Tbx1 signalling circuit associated with pharyngeal identity. Through lineage tracing, gene knockout, and inhibitor assays, we reveal that Foxa2⁺ endoderm is necessary for Tbx1 expression in the adjacent mesoderm, establishing a minimal yet mechanistically informative model of early craniofacial development.

## RESULTS

### Establishing anterior-posterior patterning through aggregate fusion

Standard mouse gastruloids lack anterior mesoderm/endoderm required for craniofacial development (Miao et al., 2023; Van Den Brink et al., 2020; Veenvliet et al., 2020). We addressed this limitation by devising a modular approach that merges two aggregates differing in axial competence at the onset of primitive streak identity (day 3 (d3) of differentiation, corresponding to ∼E6.75-7.0 embryos). EPI aggregates were generated from naïve mESCs cultured in Activin/FGF2/KSR to promote epiblast-like/anterior PS identity(Kojima et al., 2014; Brons et al., 2007; Hayashi et al., 2011; Kaufman-Francis et al., 2014; Tesar et al., 2007), whereas CHI aggregates were obtained using the standard gastruloid protocol with a CHIR99021 pulse to induce posterior/late PS fates (Van Den Brink et al., 2014) (Figure 1a). Upon juxtaposition, resulting EPICHI chimeras fused efficiently (100% within 24h; Figure S1a) and robustly elongated by d5, showing increased major axis length compared to standard (CHI) gastruloids at d5 and d6 (Figure S1b).

**Figure 1.**
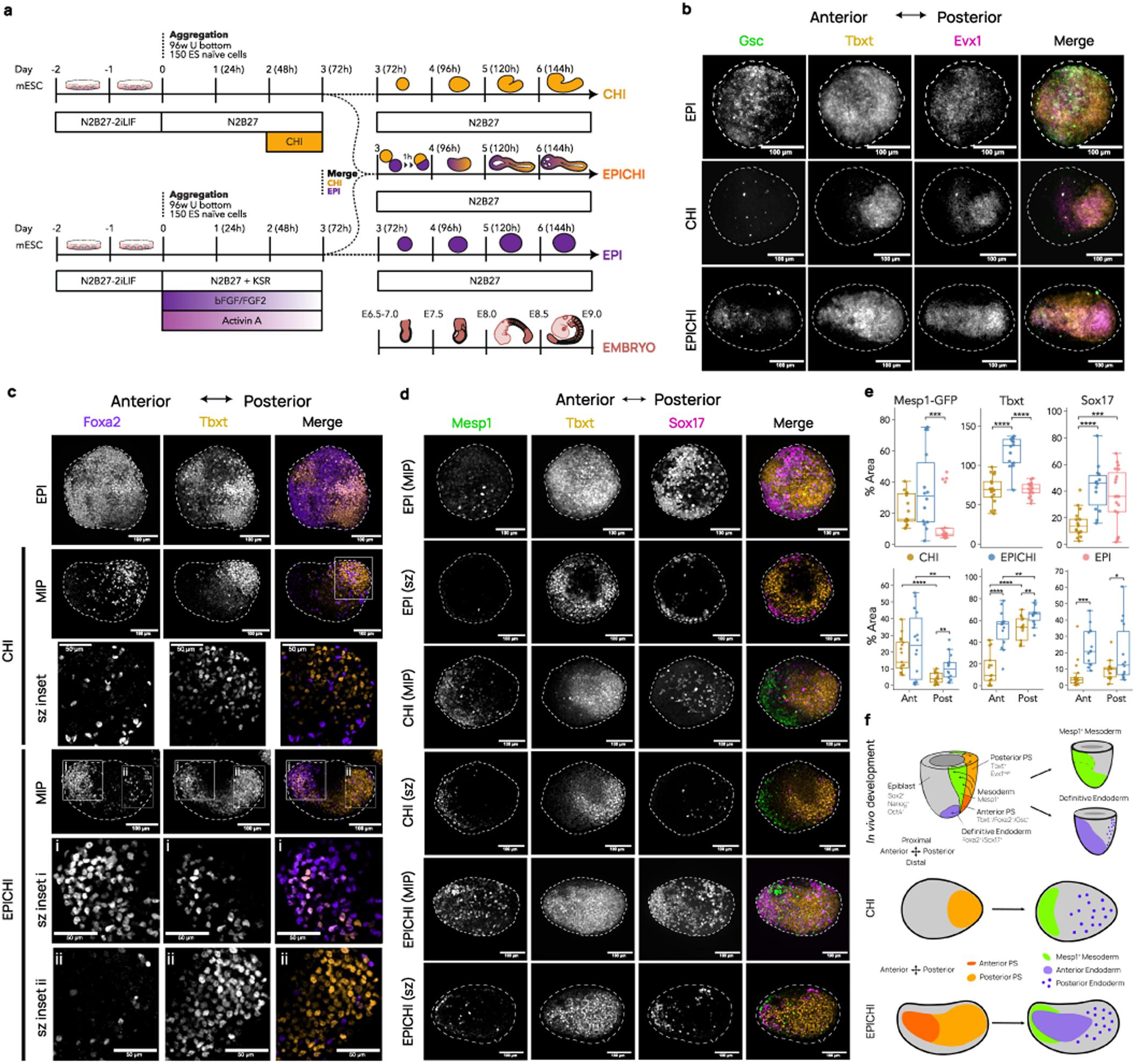
Generation of a dual-gastruloid protocol that promotes emergence of anterior and posterior primitive streak fates. **(a)** Scheme of chimera protocol. CHI, CHIR99021 **(b)** HCR staining of d4 EPI, CHI and EPICHI for *Gsc* (green), *Evx1* (magenta), and *Tbxt* (orange). A-P axis in left-right orientation. Tbxt marks posterior pole of CHI. EPICHIs display opposing poles enriched for *Gsc* (anterior) and *Evx1* (posterior). EPI lacks evident AP polarity. Scale: 100 μm. **(c)** Day 4 staining with Foxa2 and Tbxt (Brachyury) antibodies. sz inset: zoom of confocal sections of Tbxt (orange), and Foxa2 (purple) staining distributions for CHI and EPICHI (i, anterior; ii, posterior). Tbxt marks posterior pole. Scale: 100 μm (inset: 50 μm). **(d)** Day 4 staining with GFP (Mesp1) (green), Sox17 (magenta), and Tbxt (orange) antibodies. EPICHIs show anterior juxtaposition of Sox17⁺ and Mesp1⁺ cells. Scale: 100 μm. **(e)** Quantifications of positive %Area between d4 EPI, CHI and EPICHI whole samples and between anterior (Ant) and posterior (Post) halves of d4 CHIs and EPICHIs. EPI: n=19; CHI: n=17; EPICHI: n=14; 3 independent experiments/condition. **(f)** Scheme of mid-streak regionalisation in mouse embryos, CHI, and EPICHI, indicating discrete anterior/posterior domains and resulting in definitive endoderm and anterior mesoderm territories. MIP, maximum intensity projection; sz, single z-section.

Before merging, at d3, both expressed pluripotency factors Nanog/Oct4/Sox2, but only CHI expressed robustly the pan-PS marker Tbxt (Figure S1c-e). At the same stage, EPI expressed Eomes with low Tbxt and no detectable Wnt3a, whereas CHI was Tbxt-high with barely detectable Eomes and Wnt3a (Figure S1f). By d4 (mid-PS, ∼E7.5), EPI upregulated *Tbxt* along with anterior streak markers *Gsc* and Foxa2, while maintaining Nanog/Oct4 (Figure 1b-c; Figure S1c); retention of Nanog and Oct4 at this stage indicates a later entry into the primitive streak programme in EPI than in CHI. At d4, EPI expressed high levels of Tbxt and Eomes with low-Wnt3a, whereas CHI expressed Tbxt and Wnt3a with Eomes no longer detected. In EPICHIs, Eomes and Wnt3a occupied non-overlapping domains at the two ends of the fused structure while expressing Tbxt (Figure S1g). CHI showed posterior-polarized Tbxt colocalising with *Evx1* (Figure 1b-d), representing mid-to-posterior identity (Tbxt^+^/Evx1-high/Gsc-/Foxa2-) versus EPI’s mid-to-anterior identity (Tbxt+/Evx1-low/Gsc+/Foxa2+) (Figure 1b-d). EPICHIs integrated these domains within a single structure, displaying distinct Evx1-high (posterior) and Gsc+ (anterior) poles that resemble early embryonic anterior-posterior (A-P) polarity (Figure 1b). Anterior streak-specific Foxa2-Tbxt co-expression, absent in CHI, was consistently observed in EPICHI anterior regions (Figure 1c-i). Posterior Foxa2+/Tbxt-cells in CHI and EPICHI (Figure 1c-ii) likely represent posterior definitive endoderm, whereas anterior Foxa2+/Tbxt-cells in EPICHI (Figure 1c-i) likely correspond to anterior definitive endoderm. This is consistent with the localisation of Sox17⁺ definitive endoderm, robustly present in EPIs and EPICHIs close to Mesp1+ mesoderm at the EPICHI anterior interface, a configuration rarely seen in CHI gastruloids (Figure 1d). Quantifications confirm that, while Mesp1+ cells are robustly present in both CHI and EPICHI and enriched at the anterior half of the aggregates, Tbxt and Sox17 expression significantly increase in EPICHIs versus CHIs, with a major increase at the anterior pole (Figure 1e). Thus, merging two fate-biased aggregates reproducibly generates a structure with distinct anterior and posterior domains in which anterior endodermal and mesodermal progenitors are directly juxtaposed (Figure 1f).

### Spatial alignment of foregut-like endoderm and cranial-mesoderm-like cells in dual-gastruloids

From d5, EPICHIs generated a Sox17+/Foxa2+ epithelial structure (Figure 2a-b; Figure S2a-b), extending across the A-P axis (Figure 2c; Figure S2c-d), and surrounded by Mesp1-GFP⁺ mesoderm (Figure 2a-b). This epithelium co-expressed Sox2 and Sox17 anteriorly (Figure S2a), consistent with foregut-like identity in E7.75-8.0 embryos (Ikonomou and Kotton, 2015; Sherwood et al., 2009). Quantifications confirmed anterior enrichment of Foxa2+, Sox17+, Sox2+, and Tbxt+ area in EPICHIs (Figure 2c-d; Figure S2c-f; Table S1). To ask whether this Sox17⁺/Foxa2⁺ structure is an organised epithelium and whether endothelial cells contribute to it, we co-stained for Sox17, Cdh1 and Cd31. Co-staining for Sox17, Cdh1 and Cd31 showed that this structure is a Cdh1⁺ epithelium at d5 and d6 (Figure 2e). Cd31⁺ cells were detected from d5 but are adjacent to rather than within the Sox17⁺/Cdh1⁺ epithelium (Figure 2e). The anterior structure is therefore an epithelium from which Cd31⁺ cells are excluded, and not of vascular identity. In contrast, CHI aggregates displayed mainly scattered posterior Sox17+/Foxa2+ cells (Figure 2a; Figure S2a-b) with a few small, mostly posterior Cdh1⁺/Cd31⁺ cell islands (Figure 2f), while EPIs contained large Sox17+/Foxa2+/Cdh1+ areas but lacked clear A-P orientation (Figure 2f, Figure S2a-b). Occasional anterior Tbxt+/Foxa2+ cells suggested the presence of nascent notochord-like structure (∼60% (35/57) of d5 and 50% (11/22) of d6 EPICHIs) (Figure S2g; Table S2). Cells co-expressing Sox2 and Tbxt were detected in the posterior region of EPICHIs (Figure S2a), indicating that neuromesodermal-like progenitors are present in these dual-gastruloids.

**Figure 2.**
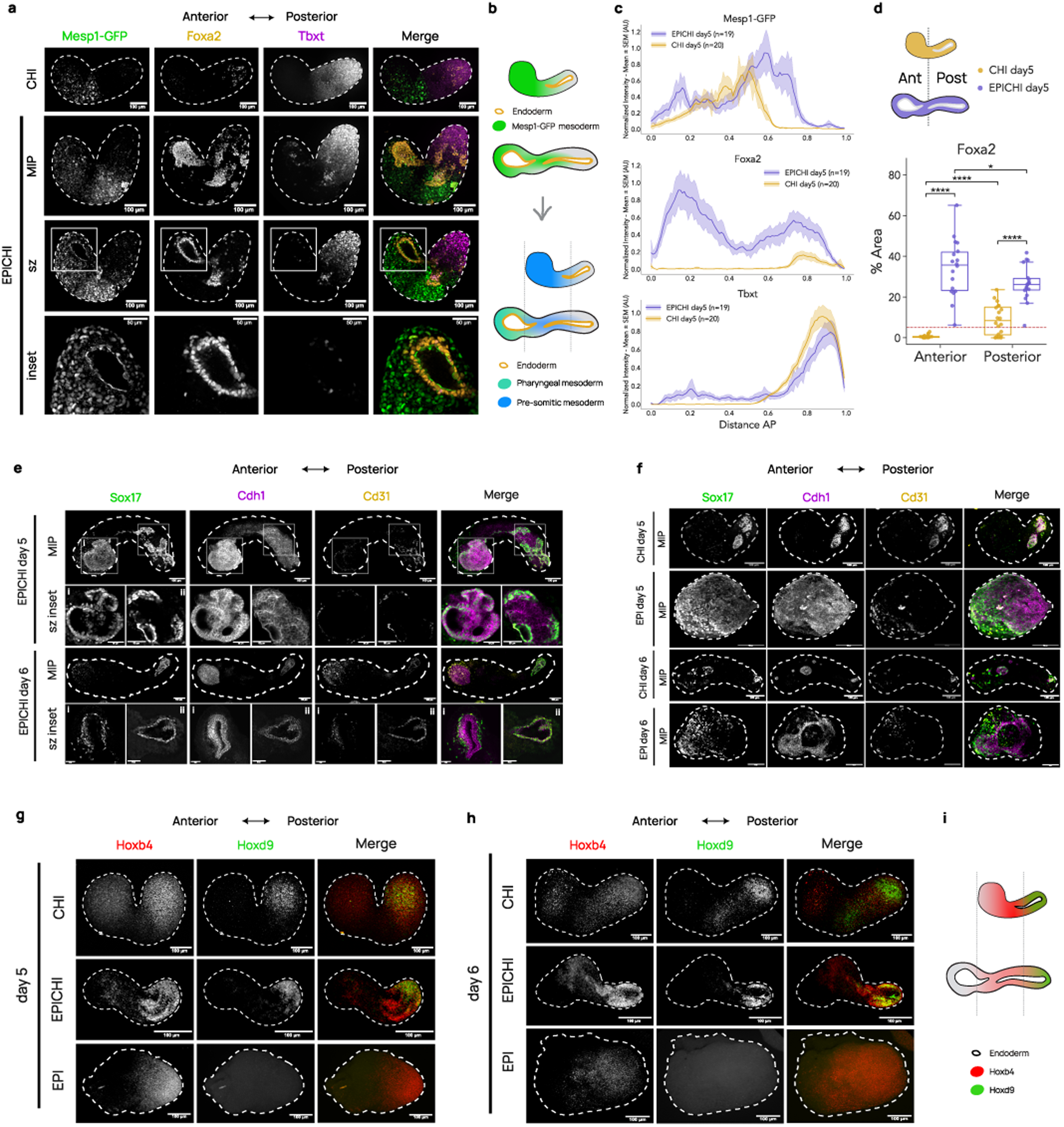
Anterior–posterior polarity and formation of a foregut-like epithelium surrounded by mesoderm in dual-gastruloids. **(a)** Day 5 Mesp1-GFP EPICHIs stained with GFP (green), Foxa2 (orange), and Tbxt (magenta) antibodies. Tbxt marks the posterior pole of CHI and EPICHI samples; Bottom: inset of confocal sections of EPICHI showing Mesp1^+^ mesoderm surrounding anterior endoderm-derived epithelial cells. MIP, maximum intensity projection; sz, confocal section. **(b)** Scheme of Mesp1-GFP expression in CHI and EPICHI, with respective presumptive cranial mesoderm and presomitic mesoderm territories surrounding endodermal epithelium. **(c)** Quantification of Mesp1-GFP, Foxa2, and Tbxt along AP axis. Bold central line shows mean value while shaded area shows SEM from multiple samples; n quantified per condition indicated in legend; N=3 independent experiments/condition. **(d)** Quantification of Foxa2 positive %Area in discrete anterior and posterior regions of d5 CHI and EPICHI. Dashed line, %Area threshold (5%) of positive and negative samples. CHI: n=20; EPICHI: n=18; 3 independent experiments/condition. **(e)** Day 5 and d6 EPICHIs stained for Sox17 (green), Cdh1 (magenta), and Cd31 (orange). sz insets: boxed anterior (i), posterior (ii) regions. Co-staining shows Sox17⁺/Cdh1⁺ epithelium, with scattered Cd31⁺ cells adjacent to, but not within, epithelium from d5. **(f)** Day 5 and d6 CHIs and EPIs stained for Sox17 (green), Cdh1 (magenta), and Cd31 (orange). A-P axis in left-right orientation for CHI. CHI samples show posterior Sox17⁺/Cd31⁺/Cdh1+ cells; EPI express epithelial Sox17^+^/Cdh1^+^ but lack an evident A-P polarity. **(g)** Day 5 EPICHIs, CHIs and EPIs stained for Hoxb4 (red) and Hoxd9 (orange). Hoxd9 is confined to posterior pole of EPICHIs and CHIs, absent from EPIs; Hoxb4 is expressed in anterior areas, overlapping with Hoxd9 and extends to anterior areas in CHI and EPICHI. **(h)** Day 6 EPICHIs, CHIs and EPIs stained for Hoxb4 (red) and Hoxd9 (orange), showing temporal maintenance of expression domains of Hoxb4 and Hoxd9. Scales: 100 μm (insets: 50 μm). **(i)** Schemes showing Hoxd9 expression at posterior-most tip in CHI and EPICHI; Hoxb4 expression from posterior to mid regions in EPICHI, and to anterior-most part in CHI.

Mesp1-GFP expression in EPICHIs peaked anteriorly (0.0-0.4 distance) and mid-to-posteriorly (0.4-0.8; Figure 2c), likely corresponding to cranial-mesoderm-like cells surrounding foregut-like endoderm (Foxa2/Sox17/Sox2 peaks at 0.0-0.4; Figure 2c; Figure S2c), and pre-somitic mesoderm (Figure 2b). In contrast, CHI samples showed a single middle peak (0.2-0.6; Figure 2c) likely representing anteriorly shifted pre-somitic mesoderm due to anterior truncation (Figure 2b). This conclusion is supported by the anterior expression of Pax1, expressed by somites, in CHI samples (Figure S4a). We next examined two Hox genes. Hoxb4 was detected in EPICHI, CHI and EPI d5 and d6; in EPICHI, Hoxb4 was detected from posterior to middle regions, while in CHI it was detected also at the most anterior tip. Hoxd9 was detected in EPICHI and in CHI, restricted to the posterior pole of the structure, and was not detected in EPI at either stage (Figure 2g-i). These results point to an extended anterior domain in EPICHI gastruloids that lacks expression of these Hox genes. Thus, EPICHIs generate a Foxa2⁺/Sox17⁺/Cdh1⁺ epithelium at the anterior pole, with Mesp1⁺ mesoderm in adjacent positions suggesting spatial alignment of anterior definitive endoderm and cranial mesoderm-like cells, and a foregut-like epithelium, a configuration comparable to the arrangement of foregut endoderm and cranial mesoderm in the E8.0–8.5 embryo.

### Lineage tracing identifies EPI as the source of foregut endoderm

To trace endodermal and mesodermal lineage contributions, we generated reciprocal EPICHIs using the constitutively expressed knock-in reporters Foxa2-Venus fusion (FVF) and Mesp1-GFP, merged with 129/SvEv wildtype aggregates (Figure 3a, d). In reciprocal configuration they resolve compartment origin for the endodermal and mesodermal lineages. Foxa2-Venus signal broadly labelled the Foxa2+ epithelium when originating from EPI (100% of EPICHIs; 13/13) but was minimal when derived from CHI (1/15 EPICHIs express it anteriorly) (Figure 3a; Figure S3a; Figure S6d-f, Tables S1-S2). A-P quantifications confirmed that Foxa2-Venus+ cells derived predominantly from EPI, particularly at the anterior region (Figure 3b,c), indicating that pharyngeal-like endoderm originates almost exclusively from EPI-derived progenitors.

**Figure 3.**
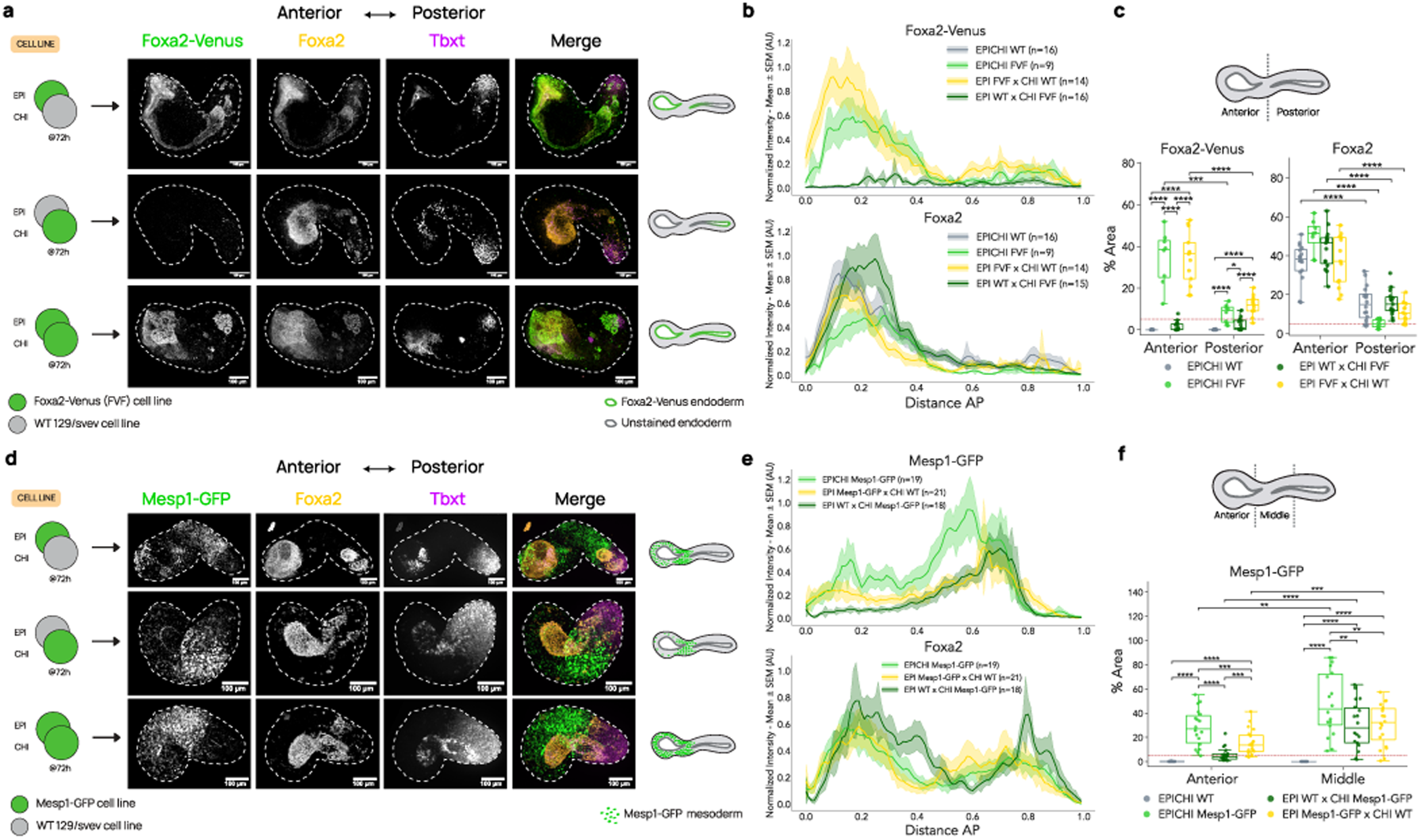
Relative contributions of EPI and CHI aggregates to endoderm and mesoderm in EPICHI reciprocal chimeras. **(a)** Design of genetic chimera protocol using Foxa2-Venus fusion protein (FVF) reporter line and wild-type 129/SvEv ES line in reciprocal combinations, merged at 72h; d5 representative images stained with GFP/Venus (green), Foxa2 (orange), and Tbxt (magenta) antibodies. Tbxt marks posterior pole of EPICHI. Right: schemes of Foxa2-Venus contribution to endodermal epithelium for each configuration. Note that endodermal Venus contribution is obtained when FVF is in EPI and is largely restricted to posterior epithelium when FVF is in CHI. **(b)** A-P quantification of GFP/Venus (Foxa2-Venus) and Foxa2 signal intensities at d5. Mean in bold; SEM from multiple samples, shaded; n quantified/condition indicated in legend. N=2 independent experiments/condition. **(c)** Quantification of GFP/Venus (Foxa2-Venus) and Foxa2 positive %Area in discrete anterior and posterior regions of d5 EPICHI chimeras. Dashed line, %Area threshold (5%) to class positive and negative samples. EPICHI WT: n=16; EPICHI FVF: n=9; EPI FVF x CHI WT: n=14; EPI WT x CHI FVF: n=15; 2 independent experiments/condition. **(d)** Design of Mesp1-GFP/wild-type 129/SvEv reciprocal chimera protocol and corresponding d5 representative images stained with GFP (green), Foxa2 (orange), and Tbxt (magenta) antibodies. Tbxt marks posterior pole of EPICHI. Right: schemes of Mesp1-GFP contribution to mesoderm. **(e)** A-P quantification of Mesp1-GFP and Foxa2 expression at d5. Mean in bold; SEM from multiple samples, shaded; n quantified/condition in legend. N=3 independent experiments/condition. **(f)** Quantification of Mesp1-GFP^+^ %Area in discrete anterior and middle regions at d5. Dashed line, %Area threshold (5%) to class positive and negative samples. EPICHI WT: n=21; EPICHI Mesp1-GFP: n=18; EPI WT x CHI Mesp1-GFP: n=18; EPI Mesp1-GFP x CHI WT: n=20; 3 independent experiments/condition. Scales: 100 μm.

Mesp1-GFP tracing revealed contributions from both aggregates (Figure 3d; Figure S3b, Figure S6a-c). EPI-derived Mesp1+ cells localized anteriorly around the Foxa2+ foregut-like epithelium (90% of EPICHIs; 18/20) and medially in presumptive presomitic regions (90% of sample; 18/20), whereas CHI-derived Mesp1⁺ cells primarily contributed to mid mesodermal territories (in ∼90% of EPICHIs; 16/18) and less to the anterior region (∼40% of EPICHIs; 7/18) (Figure 3e,f; Figure S6a-c; Tables S1-S2). Foxa2-Venus and Mesp1-GFP distribution demonstrate partial plasticity, whereby both aggregates can contribute to cranial-mesoderm-like cells, but EPI-derived cells consistently contributed to anterior positions associated with cranial-mesoderm-like and endodermal identity.

### Emergence of pharyngeal identity in endoderm and surrounding mesoderm

By d5-6, Hybridisation Chain Reaction (HCR) revealed *Pax1*/*Pax9* expression in *Foxa2*⁺ epithelium of EPICHIs, resembling E8.5 pharyngeal endoderm (Figure 4a; Figure S7c-d). A-P quantifications confirmed anterior enrichment of *Foxa2* and *Pax9* in d6 EPICHI compared with CHI (Figure 4b, c; Figure S4b). *Pax1,* expressed in both somites and pharyngeal endoderm *in vivo* (Figure 4a, top), localised slightly posterior to Pax9^+^ endoderm of d5 EPICHIs (*Pax1* peak at ∼0.3 distance, *Foxa2*/*Pax9* peak at ∼0.1), and at both pharyngeal and somitic positions by d6 (*Pax1* peaks at ∼0.2 and ∼0.4, *Foxa2*/*Pax9* peak at ∼0.2) (Figure 4a, b). The temporal expression of Pax1 and Pax9 in EPICHIs anterior endoderm matches pharyngeal endoderm maturation at E8.5-9.0 *in vivo*. In contrast, d5-6 CHI gastruloids showed anterior *Pax1* expression (0.0-0.3 distance) not associated with an organised Foxa2^+^ epithelium (Figure 4a; Figure S4a), consistent with anterior truncation and somitic identity of this anterior region of CHIs.

**Figure 4.**
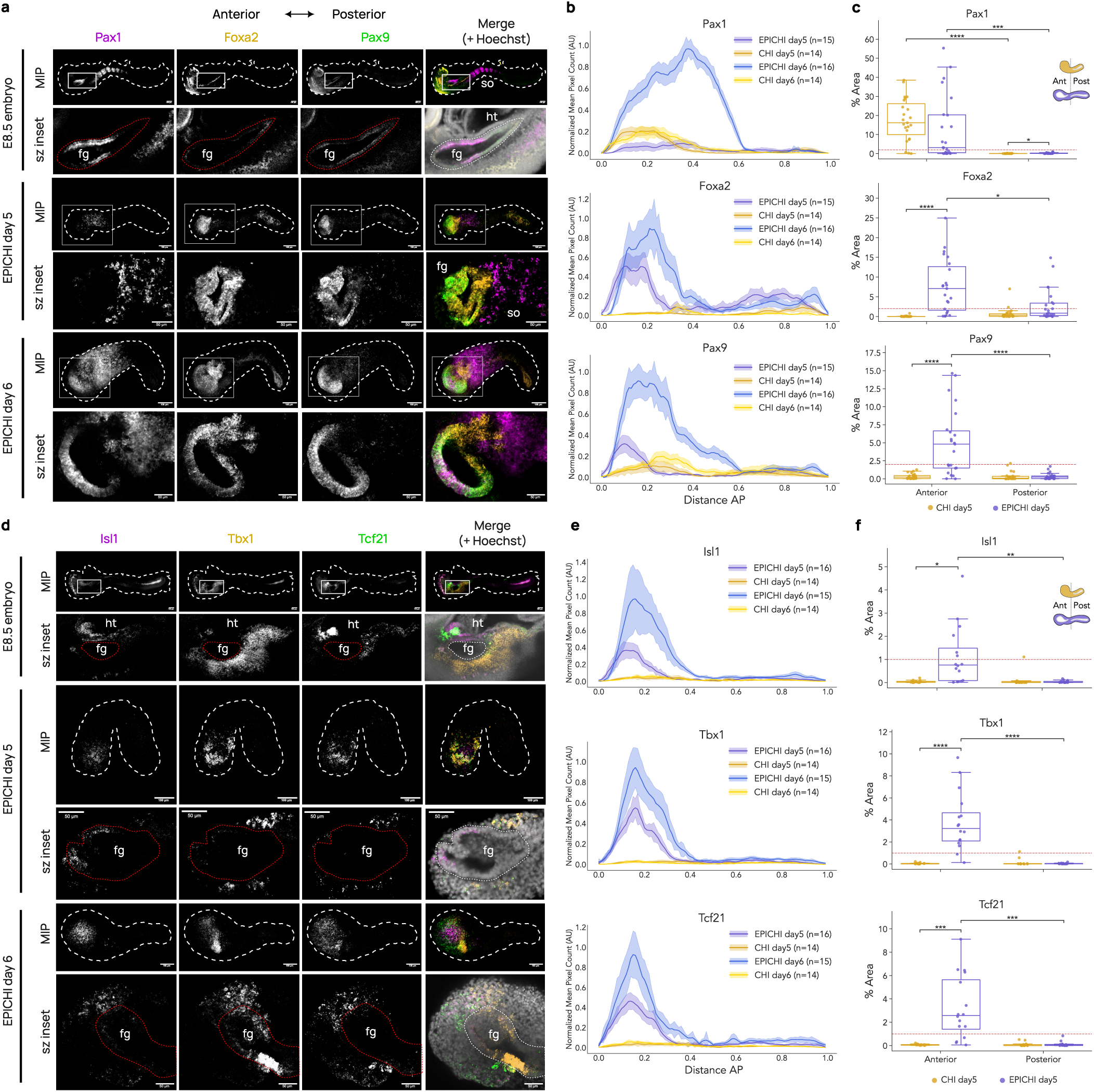
Pharyngeal-like endoderm and cranial-mesoderm-like markers in EPICHIs compared with E8.5 embryos. **(a)** E8.5 mouse embryo and d5-6 EPICHIs stained by mRNA HCR probes for *Pax9* (green), *Foxa2* (orange), and *Pax1* (magenta) merged with Hoechst (grey). Inset: detail of confocal optical sections. MIP, maximum intensity projection; sz, confocal optical section; ht, heart; fg, foregut. **(b)** Quantified A-P profiles of *Pax9*, *Foxa2*, and *Pax1* expression in CHI and EPICHI at d5 and d6. Bold central line, mean value; shaded area, SEM from multiple samples; n quantified/condition. N=2 independent experiments/condition. **(c)** Quantification of *Pax9*, *Foxa2*, and *Pax1* positive %Area in discrete anterior and posterior regions of d5 CHI and EPICHI. Dashed line, %Area threshold (2%) to class positive and negative samples. CHI: n=23; EPICHI: n=23, with 3 independent experiments/condition. **(d)** E8.5 mouse embryo and d5-6 EPICHIs stained by mRNA HCR probes for *Isl1* (magenta)*, Tbx1* (orange), and *Tcf21* (green); merge including Hoechst (grey). Inset: confocal optical sections of boxed region. ht, heart; fg, foregut. **(e)** Quantified A-P profiles for *Isl1, Tbx1,* and *Tcf21* expression in CHI and EPICHI at d5 and d6, expressed as normalised mean pixel count. Bold central line, mean value; shaded area, SEM from multiple samples; n quantified/condition. N=2 independent experiments/condition. **(f)** Quantification of *Isl1, Tbx1,* and *Tcf21* positive %Area in discrete anterior and posterior regions of d5 CHI and EPICHI. Dashed line, %Area threshold (1%). CHI: n=14; EPICHI: n=16; 2 independent experiments/condition. Scales: 100 μm (insets: 50 μm). Gastruloids: 129/SvEv cell line.

To ask whether Mesp1⁺ cells acquire cranial-mesoderm-like character, we stained for Tbx1, Tcf21 and Isl1. Tbx1 expression is common to head skeletal muscle and cardiac progenitors, while Tcf21 and Isl1 are enriched respectively in head skeletal muscle and cardiac progenitors (Alzamrooni et al., 2023; Lin et al., 2006; Moncaut et al., 2012). In EPICHIs, 93.5% (43/46) expressed Tbx1 anteriorly, 37.5% (6/16) expressed Isl1, and 75% (12/16) expressed Tcf21 (Table S2). The spatial arrangement of foregut-like endoderm surrounded by Tbx1+/Tcf21+/Isl1+ cardiopharyngeal-like mesoderm in EPICHIs resembled the configuration observed in E8.5 embryos (Figure 4d; Figure S7a-b), whereas in CHI aggregates, these genes were mostly undetectable (Figure 4e, f; Figure S4b, c). Isl1 and Tbx1 are also expressed by the foregut/pharyngeal endoderm *in vivo*; similarly, we noticed their expression in the anterior endoderm d5-6 EPICHIs (Figure 4d). EPI samples expressed both pharyngeal-like endoderm and mesoderm markers but lacked a clear A-P organisation (Figure S4a, c).

To ask whether this Tbx1⁺ domain carries a cardiac signature, we stained d5 EPICHI, CHI and EPI for *Tbx1*, *Tbx5* and *Nkx2-5*. *Tbx1* was detected in EPICHI and in EPI, whereas *Tbx5* and *Nkx2-5* were not detected in any of the three conditions (Figure S7e). Applied to E8.5 embryos, the same probe set gave *Tbx1* signal in the pharyngeal region and *Tbx5* and *Nkx2-5* signal in the cardiac region (Figure S7f).

These patterns indicate that EPICHIs combine anterior and posterior progenitors to produce foregut-like and cranial-mesoderm-like compartments resembling the E8.5 pharyngeal region, together with presomitic/somitic mesoderm, while single origin gastruloids have limited mesodermal diversity.

### Foxa2⁺ endoderm is required for cranial mesoderm-like specification

To test if endodermal signals are required for cranial-mesoderm-like identity, we generated EPICHI chimeras with Foxa2 knockout in EPI, CHI, or both aggregates (Figure 5a). Complete knockout abolished Foxa2^+^ pharyngeal-like endoderm and *Tbx1* expression in EPICHIs (Figure 5a-c) and in Foxa2-KO EPI mono-aggregates (Figure S5a). When only the EPI aggregate was composed of Foxa2-KO cells in mixed EPICHIs, we observed a phenotype comparable to the full knockout EPICHI samples lacking expression of both *Foxa2* and *Tbx1*, whereas CHI-specific knockout samples showed a full rescue in the expression of both genes (Figure 5a-c; Figure S8a-c). These results suggest that EPI-derived *Foxa2*^+^ foregut-like endoderm is required for adjacent cranial mesoderm-like specification in EPICHIs.

**Figure 5.**
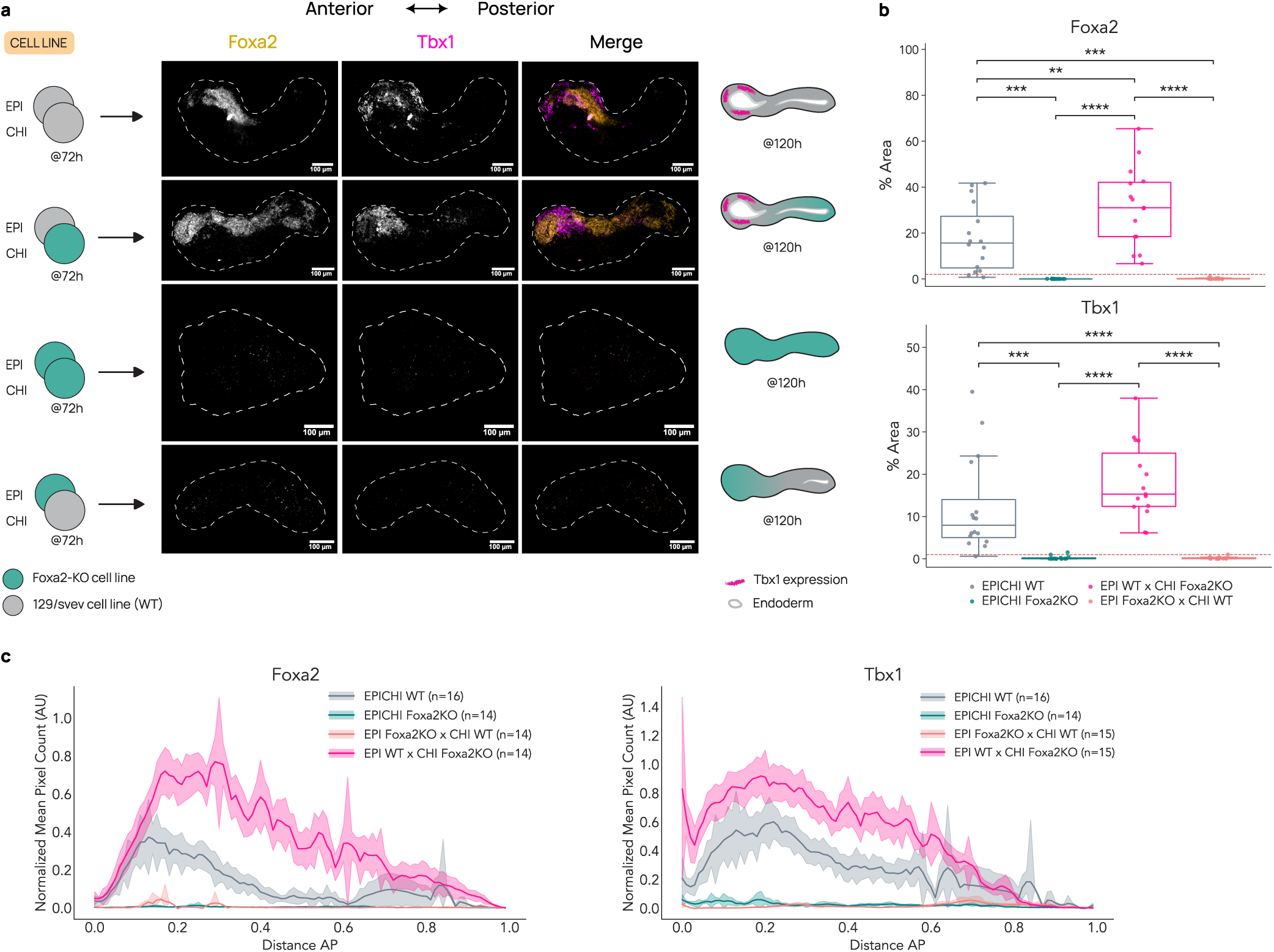
EPI-derived *Foxa2* is required for pharyngeal endoderm and *Tbx1* expression. **(a)** Design of Foxa2-KO/wild-type 129/SvEv chimeras of representative images of d5 samples stained by HCR for *Foxa2* (orange), and *Tbx1* (magenta). Scale: 100 μm. **(b)** Quantification of *Foxa2* and *Tbx1* positive %Area of d5 Foxa2-KO/wild-type EPICHI chimeras. Dashed line, %Area threshold (1-2%). EPICHI WT: n=16; EPICHI Foxa2KO: n=14; EPI WT x CHI Foxa2KO: n=15; EPI Foxa2KO x CHI WT: n=14; 2 independent experiments/condition. **(c)** AP quantification of *Foxa2* and *Tbx1* fluorescent pixels in d5 chimeras. Mean in bold; SEM from multiple samples, shaded; n quantified/condition. N=2 independent experiments.

### A *Hedgehog-Foxa2* signalling axis contributes to Tbx1 activation

To identify Tbx1-inducing cues downstream of Foxa2+ endoderm, we tested Hedgehog (Hh) signalling, known to regulate Foxa2 in pharyngeal endoderm and implicated in the development of mesoderm-derived organs (Graham and Smith, 2000; Guzzetta et al., 2020; Litingtung et al., 1998; Piotrowski and Nüsslein-Volhard, 2000; Yamagishi et al., 2003).

HCR revealed *Shh* transcript within *Foxa2^+^* pharyngeal endoderm at EPICHI anterior pole, adjacent to *Tbx1^+^* cranial mesoderm (Figure 6a; Figure S8d, e). To test whether reception of Hedgehog signal is required for this configuration, we inhibited Smoothened (Smo), the protein mediating Hedgehog signalling in receiving cells, using two distinct antagonists, Sonidegib and Vismodegib, for 24h beginning at d4, a window in which anterior endoderm and mesoderm are already present but pharyngeal identity has not yet appeared (Figure 6b). Both antagonists significantly reduced *Foxa2* and *Tbx1* by d5 (Figure 6c-e). Because the treatment window opens only after the two tissues are in place, the reduction in *Tbx1* is not attributable to a failure to generate anterior mesoderm. Together, these results identify a Hh-Foxa2-Tbx1 relationship in EPICHIs where Shh transcript is detected in the Foxa2⁺ anterior epithelium adjacent to Tbx1⁺ mesoderm, and Smoothened inhibition reduces *Foxa2* and *Tbx1* (Figure 6f).

**Figure 6.**
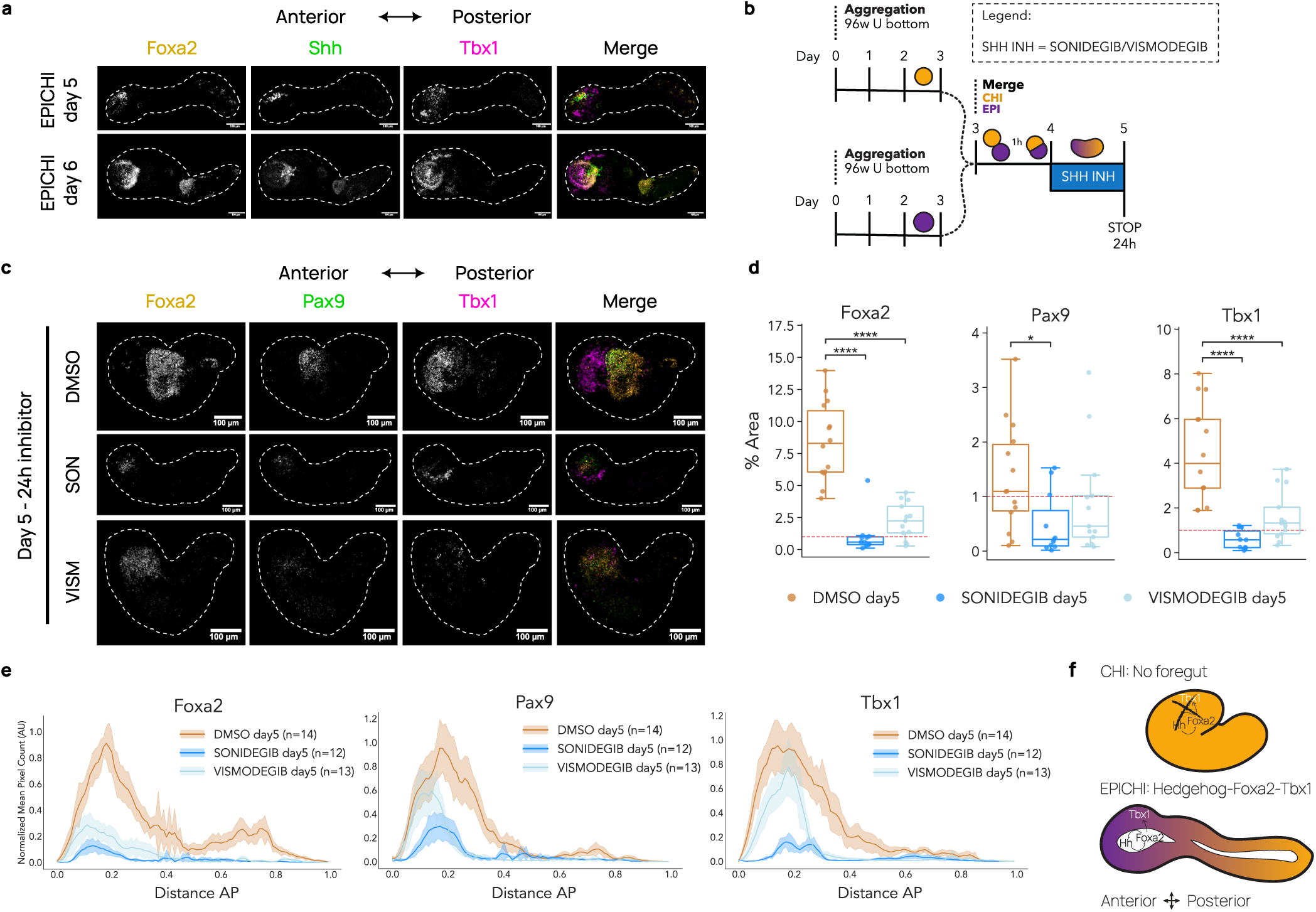
Hedgehog dependence of Foxa2+ endoderm and Tbx1 induction. **(a)** Day 5-6 EPICHIs stained by HCR for *Foxa2* (orange), *Shh* (green), and *Tbx1* (magenta). **(b)** Scheme of Smoothened inhibition in EPICHIs with 30 μM Sonidegib or 50 μM Vismodegib for 24h from 96h post-seeding (d4). **(c)** Day 5 EPICHIs after 24h treatment with inhibitors Sonidegib (SON) and Vismodegib (VISM), stained by HCR for *Foxa2* (orange), *Pax9* (green), and *Tbx1* (magenta) mRNA HCR probes. **(d)** Quantification of *Foxa2*, *Pax9*, and *Tbx1* positive %Area of d5 EPICHI after 24h of Hh inhibition with Sonidegib or Vismodegib. Dashed line represents the %Area threshold (1%). DMSO: n=14; SONIDEGIB: n=11; VISMODEGIB: n=13; 2 independent experiments/condition. **(e)** A-P quantification of EPICHIs at d5 following 24h treatment with inhibitors, respectively, stained by HCR for *Foxa2*, *Pax9*, and *Tbx1* expression. Mean in bold; SEM from multiple samples, shaded; n quantified/condition. N=2 independent experiments/condition. **(f)** Scheme of *Hedgehog-Foxa2-Tbx1* signalling axis present in EPICHI and absent in CHI. Scales: 100 μm. Gastruloids: 129/SvEv cell line.

## DISCUSSION

Stem cell-derived embryo models capture key principles of morphogenesis and lineage diversification (Arias et al., 2022; Turner and Nichols, 2023; Van Den Brink and Van Oudenaarden, 2021). Although posterior embryogenesis is well recapitulated, anterior structures including foregut, pharyngeal and cardiopharyngeal lineages essential for head muscle development remain less well represented. Here we describe a dual-aggregate gastruloid in which a foregut-like epithelium and cranial-mesoderm-like tissue arise in adjacent positions, and in which either tissue can be altered independently in chimera studies.

Our approach addresses two key limitations of current gastruloids. First, under cardiogenic conditions Mesp1+ cells give rise to first and second heart field progenitors (Rossi et al., 2021, 2022), and under cardiopharyngeal conditions gastruloids specify cardiac and skeletal muscle lineages but show no indication of an organised cranial mesoderm-like tissue (Argiro et al., 2024). Second, endodermal structures form at variable, cell-line-dependent efficiency and in several morphologies, none of which appears to have a pharyngeal endoderm-like organisation (Farag et al., 2024). EPICHIs combine two aggregates carrying different signalling histories (Activin/FGF versus CHIR pulse). Eomes and Tbxt are components of competing early and late primitive streak modules, the late module also comprising Wnt3a (Wehmeyer et al., 2025), and Nodal and Wnt/β-catenin act antagonistically in gastruloids, a declining Nodal gradient being required for anterior body structures while Wnt/β-catenin promotes posterior ones in a time– and dose-dependent manner (Dias et al., 2025). The two aggregates fall on opposite sides of this division before they meet: at d3 EPI is Eomes^+^ with low Tbxt and Wnt3a undetected, whereas by d4 CHI is Tbxt-high and Wnt3a^+^. Fusion therefore brings into contact two populations that have already entered opposing streak programmes rather than imposing a difference within a single aggregate. This configuration is associated with the co-emergence and patterning of Mesp1+ mesoderm and Sox17+/Foxa2+ endoderm into a discrete anterior compartment, and with a pharyngeal-like organisation not described in these models. Sox2^+^/Tbxt^+^ cells were also detected in the posterior region of EPICHIs, indicating that neuromesodermal-like progenitors are present in this compartment. Consistent with an extended anterior domain, Hoxb4 was detected from posterior to middle regions in EPICHI, whereas it extended to the anterior-most tip in CHI, while Hoxd9 was restricted to the posterior pole in EPICHI and CHI and was not detected in EPI.

This dual-gastruloid model results in an emerging pharyngeal-like domain in which Tbx1+/Isl1+/Tcf21+ cranial-mesoderm-like cells lie adjacent to Foxa2^+^/Pax1^+^/Pax9^+^ pharyngeal-like endoderm, resembling the early pharyngeal region of E8.0-8.5 embryos. Tbx1 and Isl1 are also expressed in cardiogenic gastruloids, where Tbx1 marks second heart field cells in populations mutually exclusive with Tbx5^+^ first heart field cells (Rossi et al., 2021); *Tbx5* and *Nkx2-5* expression were not detected in EPICHI, CHI or EPI, while the same probe set gave Tbx5 and Nkx2-5 signal in the cardiac region of E8.5 embryos. Furthermore, *Tcf21* expression is compatible with a cranial-mesoderm-like rather than a cardiac trajectory in this context. Notably, EPICHIs did not display the beating phenotype of cardiogenic gastruloids (Rossi et al., 2021, 2022). Modifying signalling cues and providing a pharyngeal endoderm-like environment therefore promotes cranial-mesoderm-like identity, and EPICHIs provide a tractable framework to uncover how lineage diversification and tissue co-development are orchestrated during early embryogenesis. Future experiments will test whether this endodermal environment actively suppresses cardiogenic potential or reinforces craniofacial identity.

Endodermal and cardiac derivatives have each been obtained in single-aggregate gastruloids. Definitive endoderm forms an endoderm-like region through cell-state transitions and collective cell movement (Hashmi et al., 2022), and develops into distinct morphotypes including a gut tube (Farag et al., 2024). Under cardiogenic conditions the cardiac domain forms next to a gut-tube-like epithelium, separated from it by an endocardial-like layer (Rossi et al., 2021, 2022), and gastruloids reproduce cardiopharyngeal mesoderm specification towards cardiac and skeletal muscle lineages, including head-like and trunk-like skeletal myoblasts (Argiro et al., 2024). In EPICHIs the anterior structure is a Sox17+/Cdh1+ epithelium from which Cd31⁺ cells are excluded, although Cd31⁺ cells are present elsewhere in the structure and in CHI aggregates. Anterior neural tissues develop in gastruloids generated without exogenous Wnt activation, in which initial aggregate size and Wnt inhibition are both crucial (Girgin et al., 2021). Combining two cell groups extends what these models reach where an engineered morphogen signalling centre acting as an organiser instructs mESC aggregates to form embryoids with a patterned primitive gut tube and anterior beating cardiac tissue (Xu et al., 2021). Chimeric gastruloids can be built by mixing two genotypes, which resolves cell-autonomous from non-autonomous gene function, or by merging two preformed aggregates, as in studies of inductive tissue interactions (Wehmeyer et al., 2025). Further, antero-posterior assembly has been combined with hypoxia in mouse (Balaskas et al., 2025), and anteriorly and posteriorly pre-patterned cells mixed at aggregation in human (Liu et al., 2025). In cardiopharyngeal gastruloids endodermal derivatives were present but may be reduced, and altered proximity to the endoderm was proposed to affect cardiac morphogenesis (Argiro et al., 2024). In EPICHIs, where the two aggregates are patterned by different signalling regimes before contact, the Foxa2-knockout chimeras and Smoothened inhibition address these questions directly (see summary Table S5).

Anterior mesoderm and definitive endoderm progenitors are among the first embryonic cell types specified during germ layer formation at the mouse primitive streak (Costello et al., 2011; Probst et al., 2021). Lineage tracing experiments with mixed *Foxa2-Venus/wt* and *Mesp1-GFP/wt* EPICHI chimeras indicate that lineage divergence likely arises during the earliest stages of mesendoderm specification in EPICHIs. Foregut endoderm arises almost exclusively from the Activin-treated aggregate, consistent with anterior definitive endoderm requiring the highest level of Nodal activity (Robertson, 2014; Vincent et al., 2003). Mesp1+ cells from both aggregates contribute to mesoderm. In most chimeras, CHI-derived Mesp1-GFP cells occupy the presomitic region, but in a subset of samples, they populate the anterior domain surrounding the foregut. This suggests that their potential can be expanded in certain conditions and supports a model in which Mesp1^+^ cells are progressively canalised toward specific mesodermal fates in response to local signalling environments (Alzamrooni et al., 2023; Chang and Kioussi, 2018; Lescroart et al., 2022). *In vivo*, subsets of Mesp1^+^ mesodermal cells that ingress through the primitive streak at proximal levels migrate towards the forming definitive endoderm (Probst et al., 2021), a region characterised by high Nodal activity. Classical transplantation experiments demonstrated that such cells preferentially contribute to cranial mesoderm (Parameswaran and Tam, 1995). Similar interactions may operate in EPICHIs, accounting for the different propensity of EPI– and CHI-derived Mesp1^+^ cells to adopt cranial fates in reciprocal chimeras. Consistent with this, *Foxa2*-knockout EPI and EPICHIs, which lack endoderm, fail to express *Tbx1* despite Activin treatment (Nodal surrogate), indicating that endoderm-derived signals are required in this context for cranial-mesoderm specification. The system thus provides experimental evidence for an early requirement of Foxa2+ foregut endoderm in establishing Tbx1^+^ cranial mesoderm. Whether *Foxa2*-deficient aggregates arrest in an undifferentiated state or divert toward alternate lineages remains to be determined.

Previous embryo studies showed that pharyngeal endoderm-secreted Shh maintains *Tbx1* expression via Foxa2 and Foxc1/2 in neighbouring endodermal and mesodermal cells (Yamagishi et al., 2003; Litingtung et al., 1998; Garg et al., 2001). In EPICHIs, *Shh* transcript is present in Foxa2^+^ pharyngeal-like endoderm adjacent to Tbx1+ cranial-mesoderm-like cells, and Smoothened inhibition reduces both Foxa2 and Tbx1. The *Hedgehog-Foxa2-Tbx1* axis examined in this study underscores the physiological relevance of EPICHIs and indicates that critical organiser-type interactions can be captured *in vitro*. This relationship, alongside possible contributions from other endodermal cues, offers one route by which foregut-like tissue influences cranial-mesoderm-like identity, and an example of how they can be manipulated pharmacologically in this system.

EPI-only aggregates express pharyngeal markers but fail to self-organise structurally in a clear anteroposterior arrangement and are biased towards mesendodermal fates, with higher Foxa2 and Foxa2/Tbxt expression, indicating that the organisation seen in EPICHIs depends on the presence of the second aggregate. The chimeric nature of EPICHIs may also provide a model to investigate morphogenetic cues required for coordinated patterning. This engineered interaction complements other modular embryo models (gastruloids (Van Den Brink et al., 2014), trunk-like structures (Veenvliet et al., 2020), cardioids (Rossi et al., 2021)) and efforts to promote anterior mesoderm identity, bridging head–trunk morphogenesis (Banerjee et al., 2025; Dias et al., 2025). Together, these platforms collectively approximate broader aspects of embryonic organisation.

While simplified, and lacking key players that shape later morphogenesis, this minimalism makes EPICHIs interpretable and allows monitoring of chimeric knockout, lineage reporter, and drug phenotypes in vitro, providing a blueprint for building higher-order region-specific embryo models. Varying the number of cells contributed by each aggregate at fusion and following intermixing directly by live imaging with a constitutive label, will determine whether the proportions set at contact constrain the extent of the anterior domain. This work extends the reach of stem-cell-based embryo models to the head, providing a conceptual and experimental framework for understanding and manipulating the earliest tissue interactions that shape craniofacial identity.

### Materials & Methods

#### Cell culture

The following mouse embryonic stem (mES) cell lines were used: 129/SvEv (EmbryoMax), Foxa2-Venus fusion (FVF) reporter line(Burtscher et al., 2013), *Foxa2^Venus/Venus^* KO (*Foxa2-KO*) line(Burtscher and Lickert, 2009), and Mesp1:p2a-NLS-dsGFP (Mesp1-GFP) reporter line. mES cells were cultured on 0.1% gelatine-coated (Sigma-Aldrich #G1890) tissue-culture plates in a humidified incubator (5% CO2, 37°C) using N2B27 medium (50% DMEM/F12 [Gibco™ #31331-093] / 50% Neurobasal [Gibco™ #21103-049] supplemented with 0.5X N2 [Gibco™ #17502-048], 0.5X B27 [Gibco™ #17504-044], 1X non-essential amino acids [Gibco™ #11140-050], 1 mM sodium pyruvate [Gibco™ #11360-070], 2 mM GlutaMAX [Gibco™ #35050-038], 1X penicillin–streptomycin [Gibco™ #15140-122], and 100 μM 2-mercaptoethanol [Gibco™ #31350-010]) supplemented with 10 ng ml−1 leukaemia inhibitory factor (Miltenyi Biotec #130-099-895), 3 μM GSK3 inhibitor CHIR 99021 (Sigma-Aldrich #SML1046) and 1 μM MEK inhibitor PDO35901 (Sigma-Aldrich #PZ0162). Cells were maintained at 20-75% confluency and passaged every 48h using TrypLE™ Express Enzyme (Gibco™ #12604-013). The passage ratios varied from 1:5 to 1:20. Cells were tested regularly for mycoplasma.

### Generation of gastruloids

CHIR99021-induced (CHI) gastruloids were generated as previously described(Van Den Brink et al., 2014). Briefly, 300 cells were seeded per well in 96-well U-bottom ultra-low attachment (ULA) plates (Corning® #7007) in 50μL N2B27 medium. After 48h, aggregates were treated with 50μL N2B27 plus 3 μM CHIR99021 (Sigma-Aldrich #SML1046) for 24h. From 72h (day 3) onward, medium was replaced daily with fresh N2B27.

EPI aggregates were generated by seeding 300 cells/well in 96-well ULA plates in 50 μL EPI medium (N2B27 supplemented with 20 ng/mL Activin A [Qkine #Qk005-0100], 12 ng/mL FGF2 [Qkine #Qk042-0100], and 1% Knockout Serum Replacement [KSR; Gibco™ #10828-010]; Hayashi et al., 2011, Girgin et al., 2021). After 48h, 50μL of N2B27 medium was added. From 72h (day 3) onward, medium was replaced daily with fresh N2B27.

To generate EPICHI aggregates, 150 cells were seeded per well and treated as above for either EPI or CHI conditions. At 72h (day 3), CHI and EPI aggregates were washed twice with N2B27 and combined in a 1:1 ratio (one aggregate each) in U-bottom ULA wells. Medium was changed daily until experiment endpoint. Cell origin was traced in reciprocal chimeras with independent reporter lines; the extent of cell mixing and any effect of fusion on fate-pool size were not quantified. Regional identities are marker– and position-based. Foxa2-Venus and Foxa2-KO EPICHIs were shorter than wild-type EPICHIs.

### Hh inhibitors treatment

EPICHI aggregates were treated with 30 μM Sonidegib (NVP-LDE225; Selleckchem #S2151) or 50 μM Vismodegib (GDC-0449; Selleckchem #S1082) for 24h beginning at 96h post-seeding (day 4).

### Embryo dissection

Mouse embryos were dissected from timed-pregnant females (B6D2F1/JRj). Embryos were separated from extraembryonic tissues in ice-cold PBS, fixed overnight in 4% paraformaldehyde (PFA) at 4°C, then dehydrated through a methanol series (25%, 50%, 75%, 100% methanol in PBS-/-containing 0.1% Tween-20 [PBST]). After storage at –20°C, samples were rehydrated through reverse methanol series and washed 3× in PBST before proceeding with staining.

### Immunofluorescence

Whole-mount immunostaining of embryo models was performed on 5-8 samples per condition. Samples were collected from 96-well plates into 1.5 mL low-bind tubes, washed 3x in PBST, fixed in 4% PFA/PBST (20 min, RT or overnight, 4°C), and stored in PBST at 4°C. After 2×5 min PBST, samples were blocked for 1h at RT in PBS containing 0.5% Triton X-100 with 10% species-matched serum and 1% BSA. Primary antibodies (Table S3) were diluted in blocking solution and incubated overnight at 4°C. Following 3×5 min PBST washes, samples were incubated with secondary antibodies in blocking solution for 4h at RT or overnight at 4°C. For goat primary antibodies, secondary antibody incubations were performed sequentially, first with donkey anti-goat, then with goat secondaries, separated by 5×5 min PBST washes to avoid cross-reactivity. After 3×5 min final washes in PBST, samples were processed for clearing/mounting (see *Clearing and mounting*).

### Hybridisation Chain Reaction (HCR)

HCR v3(Choi et al., 2018) was performed with minor modifications. Samples (5-8 per condition) were washed three times in PBS, fixed in 4% PFA/PBST (20 min RT or o/n 4°C), washed, and dehydrated through 25%, 50%, 75% methanol/PBST (5 min each) to 100% methanol, then stored in 100% methanol at –20°C for ≥1 day. Samples were rehydrated through the reverse series to PBST, followed by 1 h pre-hybridisation in hybridisation buffer (v3). Probes (0.8 µL of 1 µM stock per target in 100 µL hybridisation buffer) were incubated overnight at 37°C. Excess probe was removed by 4×15 min washes in 30% probe wash buffer at 37°C and 3×5 min washes in 5x SSCT at RT. Snap-cooled hairpins (Molecular Instruments; 1 µL of 3 µM H1 and H2 each in amplification buffer) were applied after 1h pre-incubation with amplification buffer; samples were incubated o/n in the dark at RT with gentle shaking. After washing in 5x SSCT (4×5 min, 2×30 min), samples were stained with Hoechst 33342 (4 h RT or overnight 4°C) and finally washed in PBST. Probes and hairpins are listed in Table S4.

### Clearing and mounting

Samples were gently washed with PBS under a stereomicroscope and carefully transferred onto microscopy #1.5 coverslips fitted with SS8X9-SecureSeal™ Imaging Spacers (Grace Bio-Labs GBL654008). PBS was replaced with RapiClear 1.47 clearing solution (SUNjin Lab #RC147001) before sealing with a coverslip. Imaging was performed on a Nikon Ti2E spinning-disk confocal or a BC43 CF spinning-disk confocal (Oxford Instruments) as 200-300 μm 3D z-stacks.

### Image analysis

Multichannel fluorescence images were analysed using Fiji (https://imagej.net/software/fiji/) and a custom, semi-automated Python (v3.12.9) workflow. 3D z-stacks were converted to 2D maximum intensity projections (MIPs) using Fiji. For each sample, a tissue mask was generated from the Hoechst channel and used to manually annotate a midline using Napari (v0.5.6), drawn from the anterior to the posterior side of the gastruloid, thus defining an A-P axis. The posterior pole of the axis was manually identified based on gastruloid morphology and/or Tbxt expression. Signal and background regions were manually annotated for each channel. Intensity thresholds were determined via either two-components Gaussian mixture model (GMM) with the threshold set at the midpoint between the means of the two fitted components, or the P99 method (threshold set at the 99th percentile of annotated background pixel intensities). Signal quantification was performed in a midline-based coordinate system where position 0 represents the anterior end and 1 the posterior end of the gastruloid. Each Signal-positive pixels (above the defined threshold) were assigned to its nearest midline pixel using Euclidean Distance Transform. Signal-positive pixels were binned into 100 equidistant segments along the midline, with summed intensity and pixel counts recorded per bin. All pixel values were normalized with the background threshold (the applied threshold was subtracted from each pixel value before summation). All analyses used scikit-image, scipy, seaborn, pandas, and NumPy. Samples with shape or signal artefacts that prevented reliable quantification, as assessed by visual inspection, were excluded from the analysis. EPI samples failed to elongate and were excluded from analysis due to lack of a clear anteroposterior axis.

Spatial expression profiles were analysed using a custom Python pipeline. For each condition, binned intensity data from replicate samples were aggregated and normalized to the maximum value of the condition. Condition-averaged profiles were smoothed using a Savitzky-Golay filter (window length=7, polynomial order=2) and visualised as mean ± standard error of the mean (SEM). All visualisations were generated with Matplotlib.

Measurements of the fractional area (%Area) of discrete anatomical regions (anterior, middle and posterior) were performed with a custom JavaScript macro running in Fiji (ImageJ 1.x). The user draws a line spanning the structure, starting at the anterior pole; the macro resamples it to a uniform spacing (0.5 px), computes the cumulative arc-length and extracts the midpoint (for a two-region split) or the two 1⁄3-points (for a three-region split). Adjustable perpendicular vectors are drawn through these points. Total AP line length is saved in a table and used for downstream comparisons of mid-line length between CHI and EPICHI conditions. A closed contour is drawn around the entire structure. The contour is intersected with the perpendicular line(s); the intersection points are inserted into the contour to automatically split it into two or three closed ROIs (anterior, middle, posterior). All ROIs are added to the ROI Manager and saved as a zip file. For each fluorescence channel the user sets an intensity threshold. “Analyze Particles…” is run on the calibrated channel image with analysis limited to each previously defined ROI. For every ROI–channel the macro records calibrated ROI area, %Area (Σ particle area / ROI area × 100), mean intensity, integrated density, circularity and solidity. %Area and mid-line length were compared using boxplots overlaid with stripplots of the individual measurements. Samples with shape or signal artefacts that prevented reliable quantification, as assessed by visual inspection, were excluded from the analysis. All plots were generated with Matplotlib.

### Statistical analysis

Statistical comparisons were performed on %Area using ordinary least squares regression with biological replicate included as a fixed effect. For each region, differences between conditions were tested using the model: %Area ∼ replicate + condition. For each condition, differences between regions were tested using the model: %Area ∼ replicate + region. Pairwise comparisons were conducted using two-sided t-tests on model coefficients: comparisons against the reference level test on a single coefficient, and comparisons between two non-reference levels test the difference of the corresponding coefficients. P-values were adjusted across all pairwise comparisons within each marker using the Holm-Bonferroni family-wise error rate (FWER)-controlling correction method (α = 0.05). Comparisons that were not significant after adjustment are not annotated on the plots. Statistical analyses were computed using statsmodels v0.14.5 and statistical annotations were applied to the plots using the statannotations package.

For percentage positivity calculations, marker-specific thresholds were applied based on the following criteria: 5% positive area for immunofluorescence (IF) of Sox17, Foxa2, Foxa2-Venus, and Mesp1-GFP (to select for organized cell populations) and 2% for IF of Tbxt (to select also sparse anterior notochordal-like cell streaks); 1–2% positive area for hybridisation chain reaction (HCR) staining of Pax9, Pax1, Foxa2, Tcf21, Tbx1, and Isl1 (accounting for punctate mRNA signal occupying less area than nuclear protein staining).

## RESOURCE AVAILABILITY

### Lead contact

Requests for further information and resources should be directed to lead contact, Shahragim Tajbakhsh.

### Materials availability

Mouse embryonic stem cell lines used are available with a completed materials transfer agreement. This study did not generate new cell lines.

### Data and code availability

Microscopy data will be shared by lead contact upon request. Original code is deposited at GitHub and is publicly available as of publication date; DOI is listed in key resources table. Additional information required to reanalyse data reported is available from lead contact upon request.

## Acknowledgements

We acknowledge funding support from Institut Pasteur, Agence Nationale de la Recherche (Laboratoire d’Excellence REVIVE, Investissement d’Avenir; ANR-10-LABX-73), ERC Advanced Grant (ST, 101055234). We gratefully acknowledge the UtechS Photonic BioImaging (Imagopole), C2RT, Institut Pasteur, supported by the French National Research Agency (France BioImaging; ANR-10–INSB–04; Investments for Future) for support. We are grateful to Hannah Turton and Marie Oury for help in initial stages of this project, Thomas Gregor, Isma Bennabi, Judith Pineau, Denis Duboule and Hocine Rekaik for advice and reagents, Alfonso Martinez Arias, Ulla-Maj Fiuza, André Dias for valuable discussions and advice.

## Author contributions

GM: Conceptualisation, Methodology, Software, Formal analysis, Data Curation, Investigation, Writing – original draft, Writing – review & editing, Visualisation, Supervision; MT: Software, Data Curation, Investigation, Writing – review & editing; PD: Investigation, Data Curation, Writing – review & editing; FM: Software; AM, IB, and HL: Resources; ST: Conceptualisation, Writing – original draft, Writing – review & editing, Supervision, Funding acquisition.

## Supplemental information

Figures S1–S8

Table S1. Statistical tests results

Table S2. Percentage Area over threshold

Table S3. List of antibodies used in this study

Table S4. List of HCR probes and hairpins used in this study

Table S5. Comparison with other gastruloid models

## FIGURE LEGENDS

**Figure S1.**
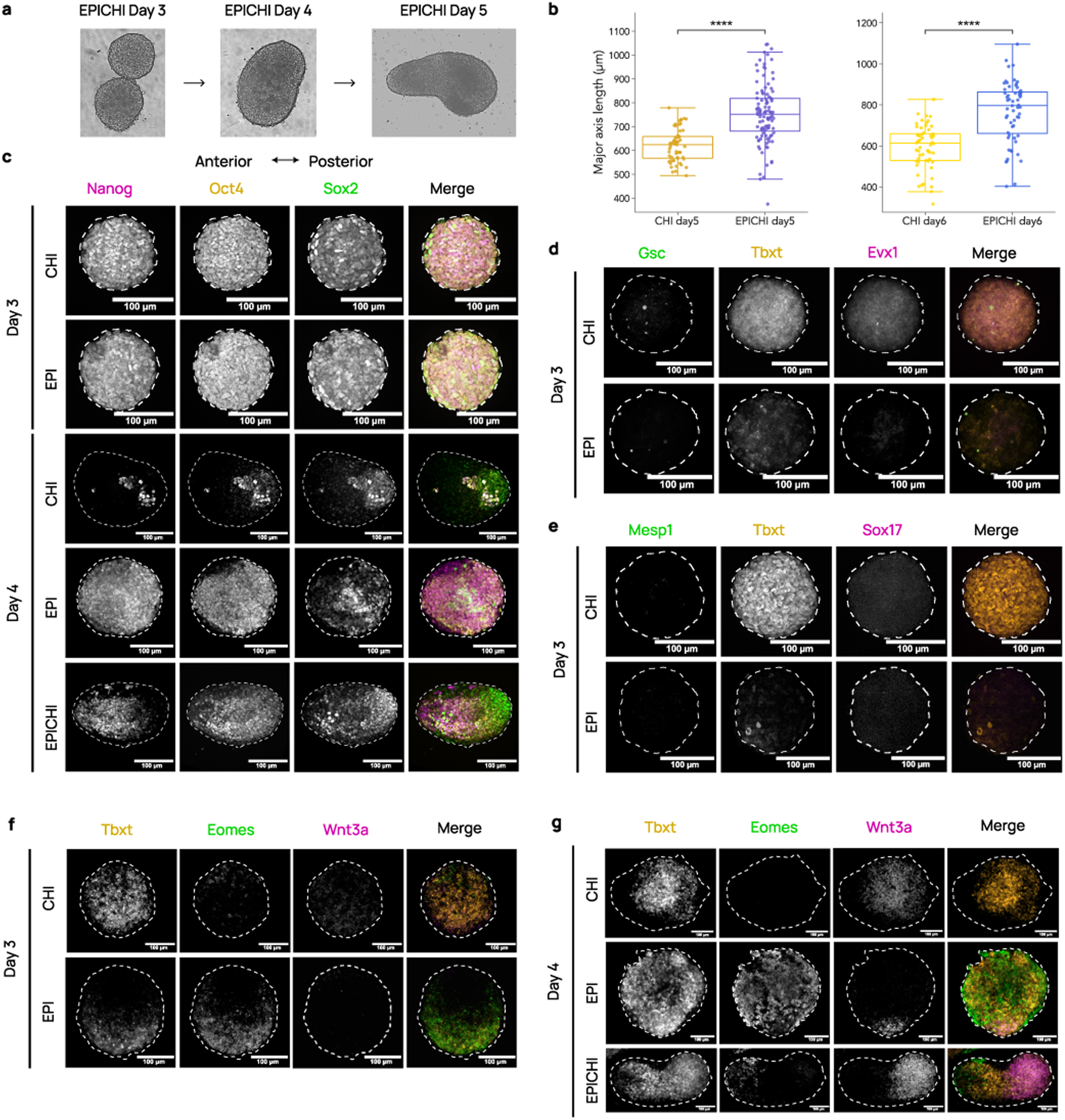
EPICHI chimeric structures fuse efficiently, elongate, and progressively exit pluripotency, and EPI and CHI adopt anterior and posterior primitive streak identities. **(a)** Bright-field microscope images of EPICHIs at d3, just before merging of EPI and CHI aggregates, at d4 24h after merging, and at d5. **(b)** Major axis length (μm) of EPICHI versus CHI samples at d5 and d6. CHI d5: n=58; EPICHI d5: n=107; CHI d6: n=56; EPICHI d6: n=60. **(c)** Day 3 and 4 EPIs, CHIs and EPICHIs stained for pluripotency markers Nanog (magenta), Oct4 (orange), and Sox2 (green). **(d)** Day 3 EPI and CHI aggregates stained with mRNA HCR probes for anterior, posterior, and pan primitive streak markers *Gsc* (green), *Evx1* (magenta), and *Tbxt* (orange), respectively. **(e)** Day 3 EPI and CHI aggregates stained for mesodermal (*Mesp1*, green; *Tbxt*, orange) and endodermal (Sox17, magenta) markers. **(f)** Day 3 EPI and CHI aggregates stained with mRNA HCR probes for *Tbxt* (orange), *Eomes* (green), and *Wnt3a* (magenta). CHI aggregates expressed *Tbxt* broadly and lacked Eomes, whereas EPI aggregates showed low *Tbxt* with detectable *Eomes*; *Wnt3a* was absent from both conditions. **(g)** Day 4 EPI, CHI and EPICHI aggregates stained with mRNA HCR probes for *Tbxt* (orange), *Eomes* (green), and *Wnt3a* (magenta). By d4, CHI aggregates were *Tbxt*⁺/*Wnt3a*⁺ and *Eomes*⁻, whereas EPI aggregates upregulated *Tbxt* and were *Tbxt*⁺/*Eomes*⁺ with a restricted *Wnt3a*⁺ domain, indicating a delayed and *Eomes*-associated onset of *Tbxt* in EPI compared with CHI. EPICHIs had both signatures within a single structure, with *Wnt3a* confined to the CHI-derived posterior domain. Scales: 100 μm.

**Figure S2.**
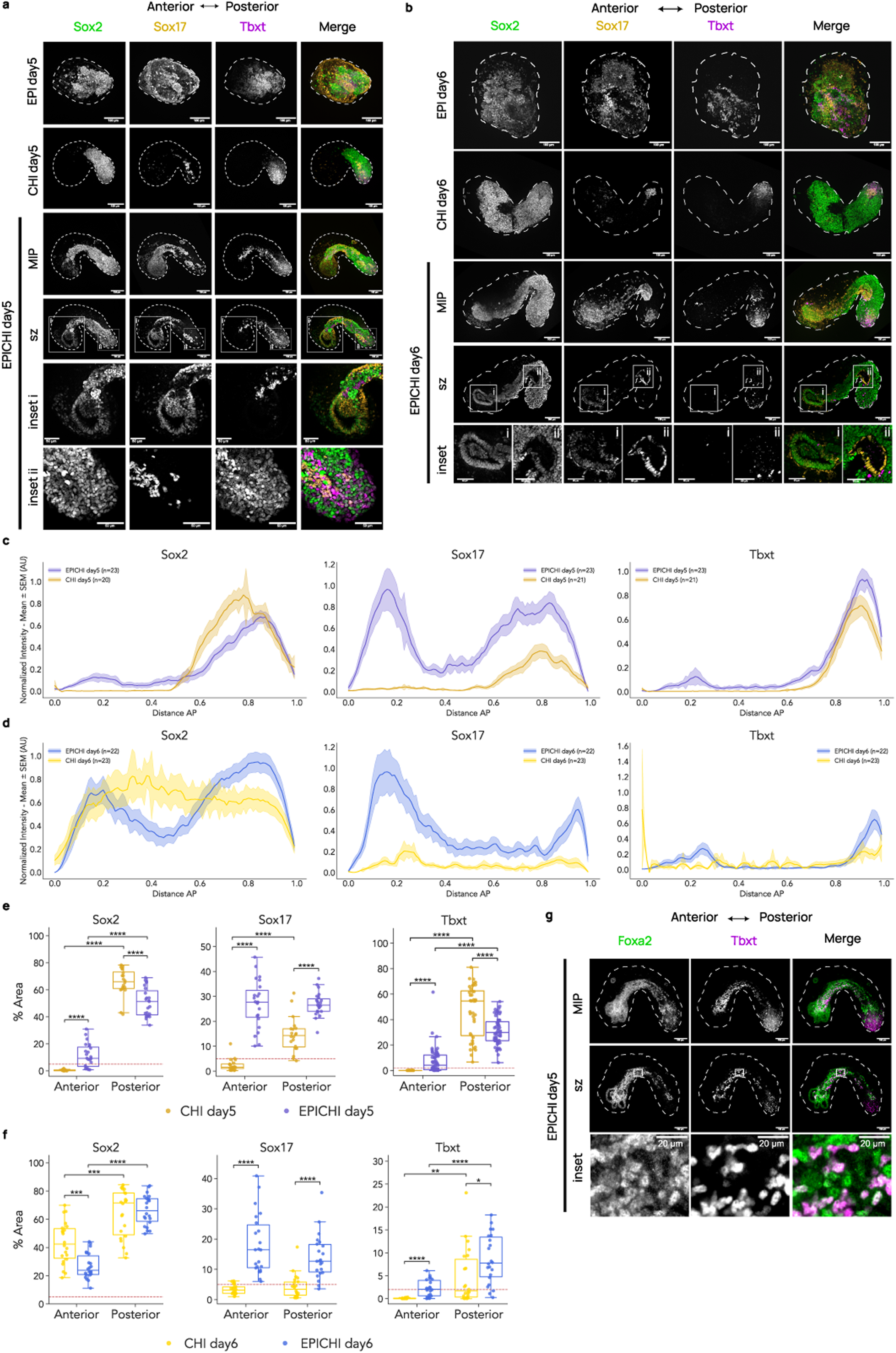
Formation of a Sox17⁺/Sox2⁺ foregut-like epithelium, posterior Sox2⁺/Tbxt⁺ neuromesodermal-like cells, and Foxa2⁺/Tbxt⁺ notochordal-like cells in EPICHIs. **(a)** Day 5 EPIs, CHIs and EPICHIs stained for Sox2 (green), Sox17 (orange), and *Tbxt* (magenta). A-P axis in left-right orientation. *Tbxt* marked the posterior pole of CHI and EPICHI samples; EPI samples lacked an evident A-P polarity. Bottom rows: confocal sections and insets of boxed anterior (i) and posterior (ii) regions of EPICHI. Inset i shows Sox17⁺/Sox2⁺ foregut-like epithelium; inset ii shows Sox2⁺/Tbxt⁺ co-expressing cells in posterior region, compatible with neuromesodermal progenitors. Scale: 100 μm (insets: 50 μm). **(b)** Day 6 EPIs, CHIs and EPICHIs stained for Sox2 (green), Sox17 (orange), and Tbxt (magenta). Tbxt marks posterior pole of CHI and EPICHI samples; EPI samples lack an evident A-P polarity. Bottom rows: confocal sections and insets showing a high magnification of foregut (i) and mid/hindgut (ii). Foregut shows higher Sox2/Sox17 expression ratio; mid/hindgut shows lower Sox2/Sox17 ratio. Scale: 100 μm (inset: 50 μm). **(c)** Quantification of A-P expression profiles for Sox2, Sox17, and *Tbxt* in d5 CHI and EPICHI. Bold central line shows mean value; shaded area shows SEM from multiple samples; n quantified per condition indicated in legend. N=3 independent experiments/condition. **(d)** A-P quantification of Sox2, Sox17 and *Tbxt* expression in d6 CHI and EPICHI. Mean in bold; SEM from multiple samples, shaded; n quantified/condition indicated in legend. N=3 independent experiments/condition. **(e)** Quantification of Sox2, Sox17 and *Tbxt* positive %Area in discrete anterior and posterior regions of d5 CHI and EPICHI. Dashed line, %Area threshold (2-5%) to classify positive and negative samples. For Sox2 and Sox17, CHI: n=20; EPICHI: n=23; 3 independent experiments/condition; for *Tbxt*, EPICHI: n=57; 8 independent experiments, CHI: n=40; 6 independent experiments. **(f)** Quantification of Sox2, Sox17 and *Tbxt* positive %Area in discrete anterior and posterior regions of d6 CHI and EPICHI. Dashed line, %Area threshold (2-5%) to classify positive and negative samples. CHI: n=22; EPICHI: n=22 3 independent experiments/condition. **(g)** Day 5 EPICHIs show colocalisation of Foxa2 (green) and *Tbxt* (magenta) in a streak of Foxa2⁺/Tbxt⁺ cells converging at midline and extending from *Tbxt*⁺ posterior tip to anterior pole, compatible with presumptive notochordal cells. MIP, maximum intensity projection; sz, confocal section. Scale: 100 μm (inset: 20 μm).

**Figure S3.**
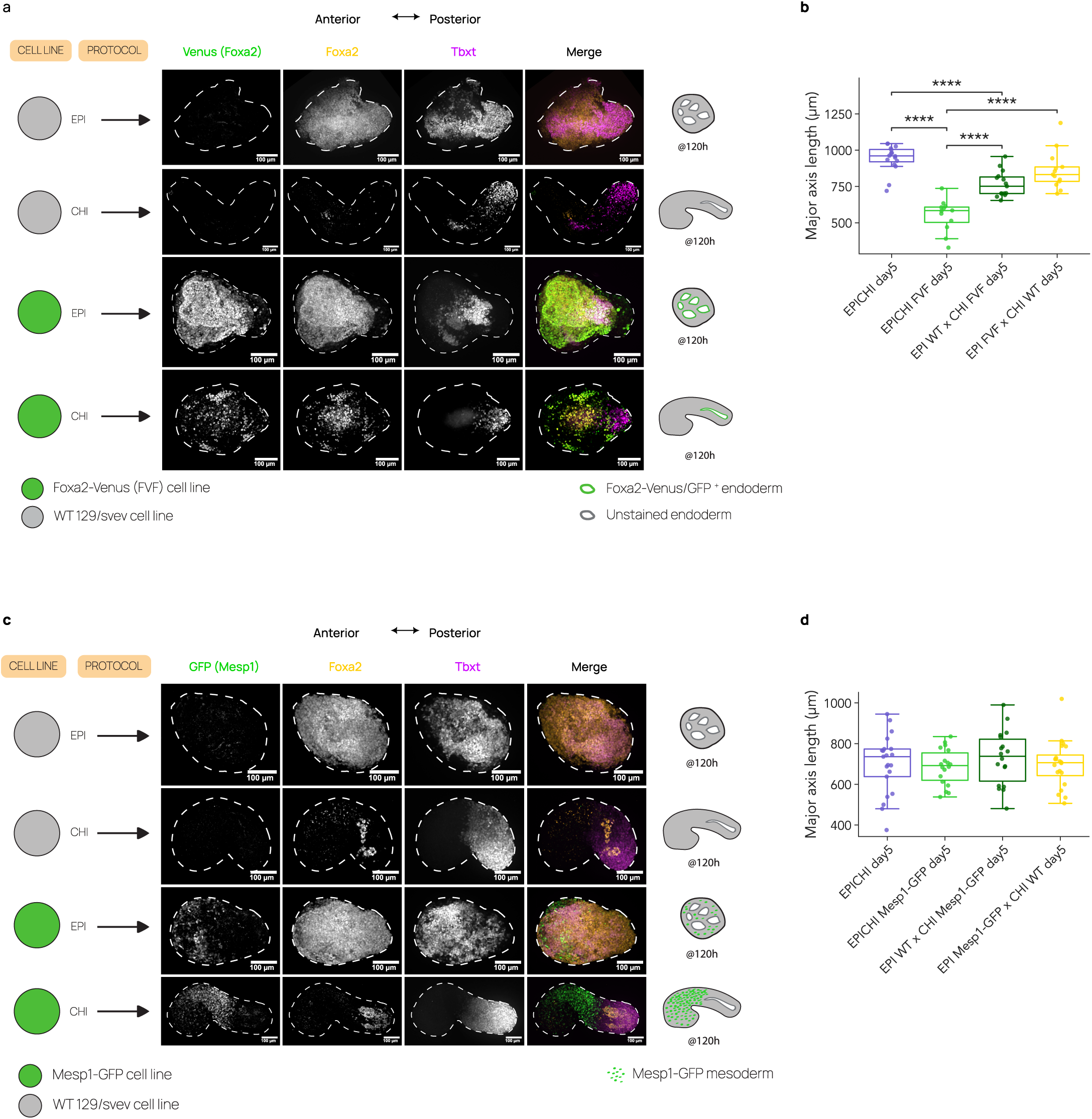
Foxa2 and Mesp1 distribution in Foxa2-Venus and Mesp1-GFP EPI and CHI mono-aggregates, and elongation of reporter-line chimeras. **(a)** Design of the generation of EPI and CHI samples using Foxa2-Venus fusion protein (FVF) reporter line and wild-type 129/SvEv cell lines, and d5 representative samples stained with GFP/Venus (green), Foxa2 (orange), and Tbxt (magenta) antibodies. A-P axis in left-right orientation. Tbxt marked the posterior pole of CHI. EPI samples lacked a clear A-P organisation and broadly expressed Foxa2, with Tbxt expression not confined to one pole and colocalising with Foxa2. Right: schemes of endodermal Venus contribution for each condition. Scale: 100 μm. **(b)** Major axis length (μm) of d5 EPICHI WT, EPICHI FVF, and the two reciprocal FVF chimeras. **(c)** Design of the generation of EPI and CHI samples using Mesp1-GFP reporter line and wild-type 129/SvEv cell lines, and d5 representative samples stained with GFP (green), Foxa2 (orange), and Tbxt (magenta) antibodies. Tbxt marked the posterior pole of CHI. EPI samples lacked a clear A-P organisation and broadly expressed Foxa2, with Tbxt expression not confined to one pole and colocalising with Foxa2. Right: schemes of Mesp1-GFP mesodermal contribution for each condition. Scale: 100 μm. **(d)** Major axis length (μm) of d5 EPICHI WT, EPICHI Mesp1-GFP, and the two reciprocal Mesp1-GFP chimeras, showing no significant difference between configurations.

**Figure S4.**
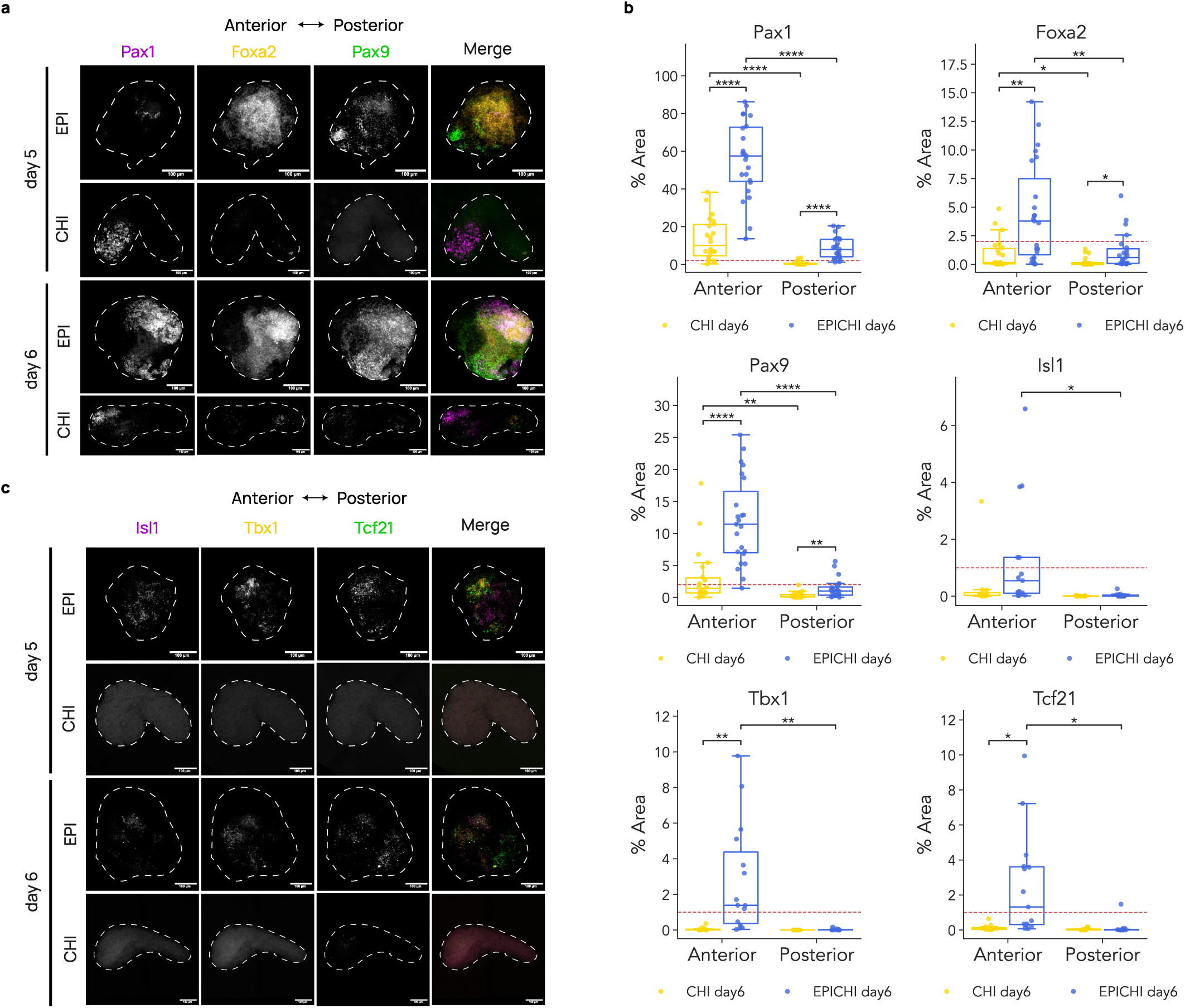
Expression of pharyngeal mesoderm and endoderm markers in EPI and CHI monoaggregates. **(a)** Day 5-6 EPI and CHI aggregates stained with mRNA HCR probes for *Pax9* (green), *Foxa2* (orange), and *Pax1* (magenta). A-P axis for CHI samples in left-right orientation. EPI samples lack a clear A-P organisation and broadly expressed *Foxa2*, with *Pax1* and *Pax9* expression increasing from d5-6. CHI samples mostly lacked expression of *Foxa2* and *Pax9* and showed anterior *Pax1* expression, compatible with somitic identity. Quantifications for d5 CHI shown in Figure 4. **(b)** Quantification of *Pax9*, *Foxa2*, *Pax1*, *Isl1*, *Tbx1*, and *Tcf21* positive %Area in discrete anterior and posterior regions of d6 CHI and EPICHI. Dashed line, %Area threshold (1-2%) to classify positive and negative samples. CHI: n=22; EPICHI: n=23; 3 independent experiments/condition for *Pax9*, *Foxa2*, *Pax1*; CHI: n=12; EPICHI: n=15; 2 independent experiments/condition for *Isl1*, *Tbx1*, *Tcf21*. **(c)** Day 5-6 EPI and CHI aggregates stained with mRNA HCR probes for *Tcf21* (green), *Tbx1* (orange), and *Isl1* (magenta). A-P axis for CHI samples in left-right orientation. EPI samples showed expression of *Isl1*, *Tbx1*, and *Tcf21* but lacked a clear AP organisation. CHI samples mostly lacked expression of *Isl1*, *Tbx1*, and *Tcf21* expression (quantifications shown in Figure 4). Scales: 100 μm

**Figure S5.**
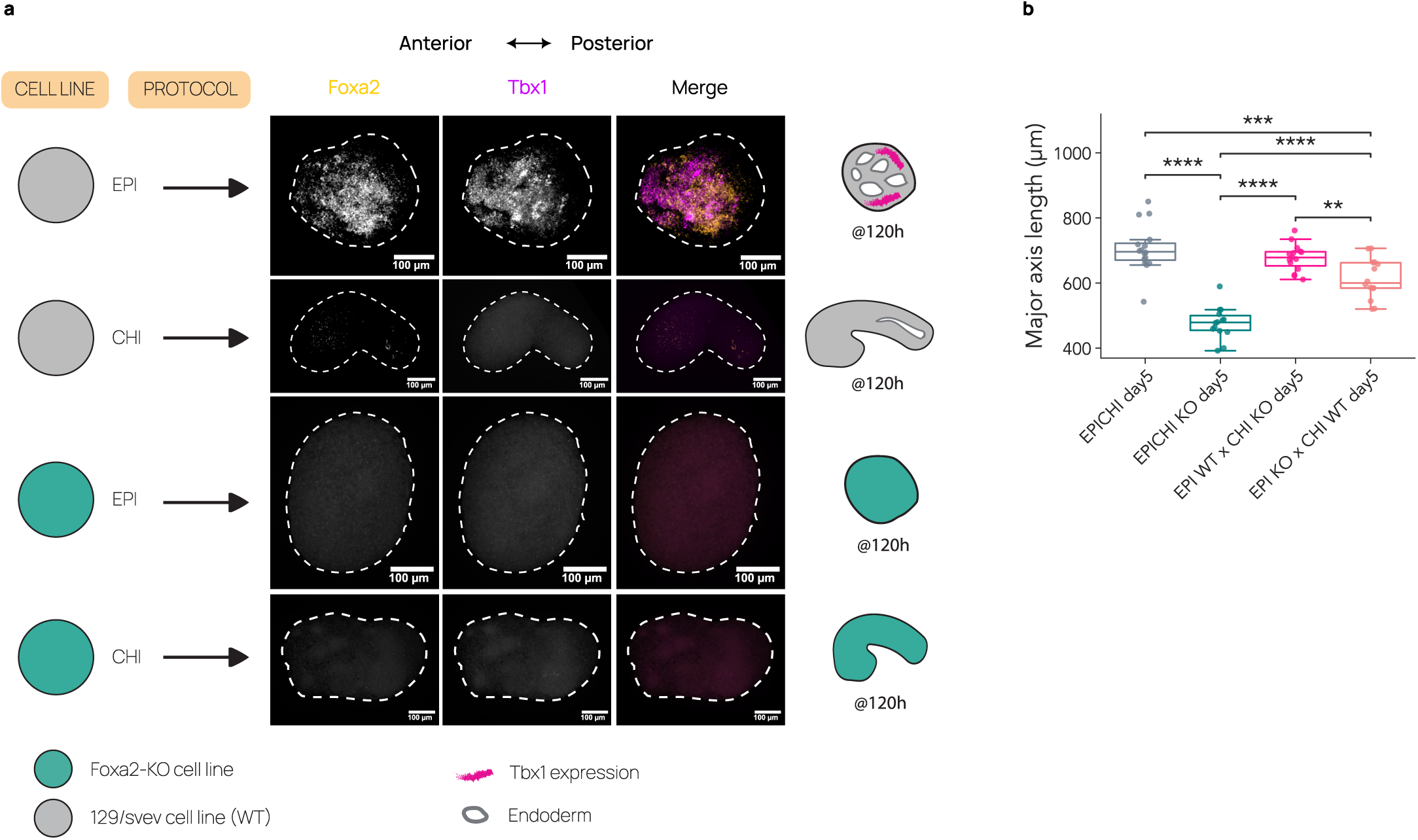
Expression of *Tbx1* in Foxa2-KO EPI and CHI mono-aggregates. **(a)** Design of Foxa2-KO/wild-type 129svev EPI and CHI aggregates with representative images of corresponding d5 EPI and CHI aggregates stained with HCR probes for *Foxa2* (orange), and *Tbx1* (magenta). A-P axis in left-right orientation. EPI samples lacked a clear AP organisation. Scale: 100 μm **(b)** Major axis length (μm) of d5 EPICHI WT, EPICHI Foxa2-KO, and the two reciprocal Foxa2-KO chimeras.

**Figure S6.**
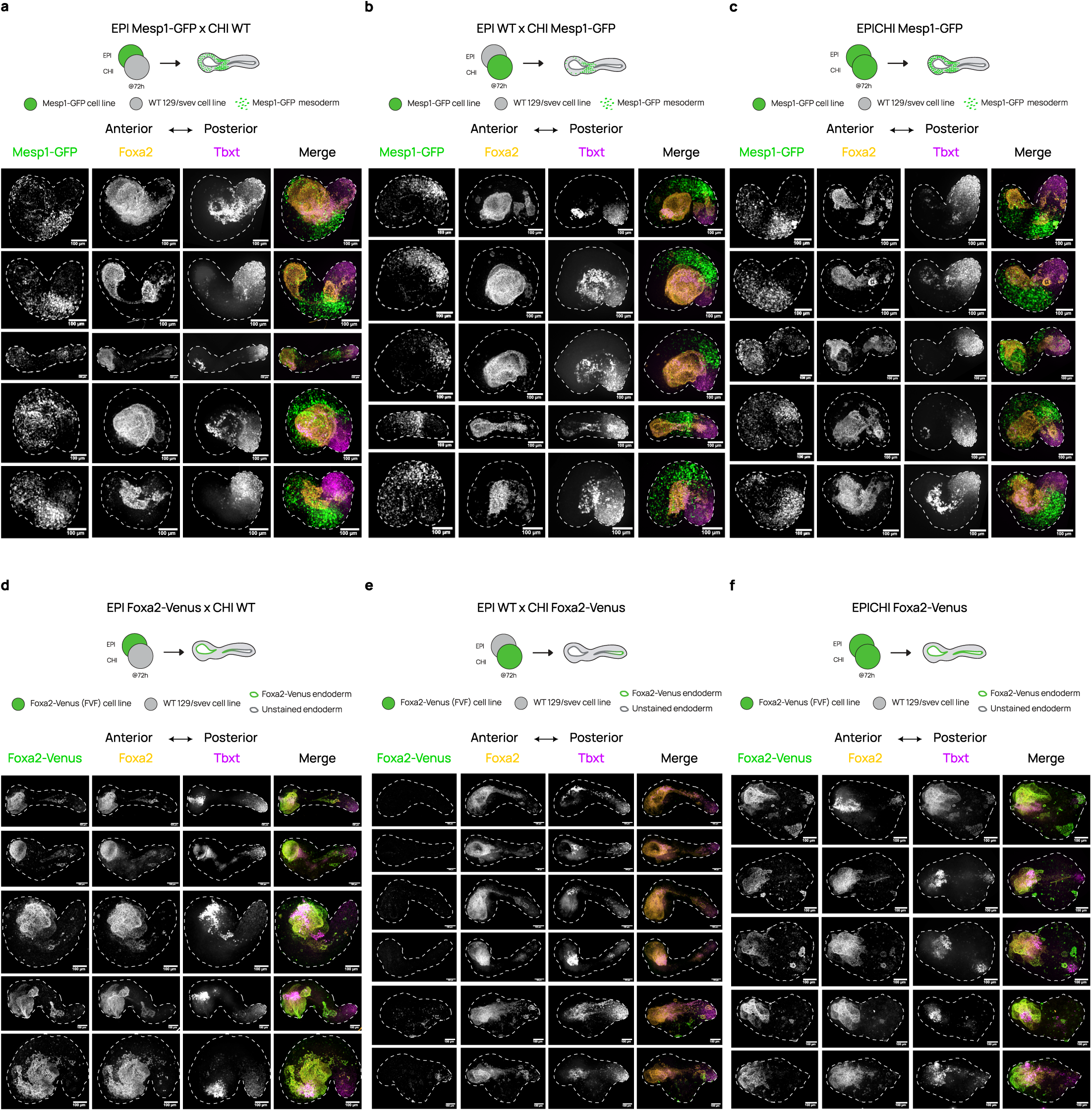
Reproducibility of EPI and CHI contributions to mesoderm and endoderm from dual-aggregate gastruloids. (**a-c**) Day 5 Mesp1-GFP reciprocal chimeras stained with GFP (green), Foxa2 (orange), and Tbxt (magenta) antibodies: (a) EPI Mesp1-GFP x CHI WT, (b) EPI WT x CHI Mesp1-GFP, (c) EPICHI Mesp1-GFP. Five independent samples are shown per configuration, with the corresponding merging scheme above each panel. A-P axis in left-right orientation. Labelled mesoderm arises from both aggregates in all configurations. **(d-f)** Day 5 Foxa2-Venus reciprocal chimeras stained with GFP/Venus (green), Foxa2 (orange), and Tbxt (magenta) antibodies: (d) EPI Foxa2-Venus x CHI WT, (e) EPI WT x CHI Foxa2-Venus, (f) EPICHI Foxa2-Venus. Five to six independent samples are shown per configuration, with the corresponding merging scheme above each panel. Venus⁺ anterior endoderm was consistently obtained when the FVF line was placed in EPI (d, f) and was largely absent when it is placed in CHI (e). Scales: 100 μm.

**Figure S7.**
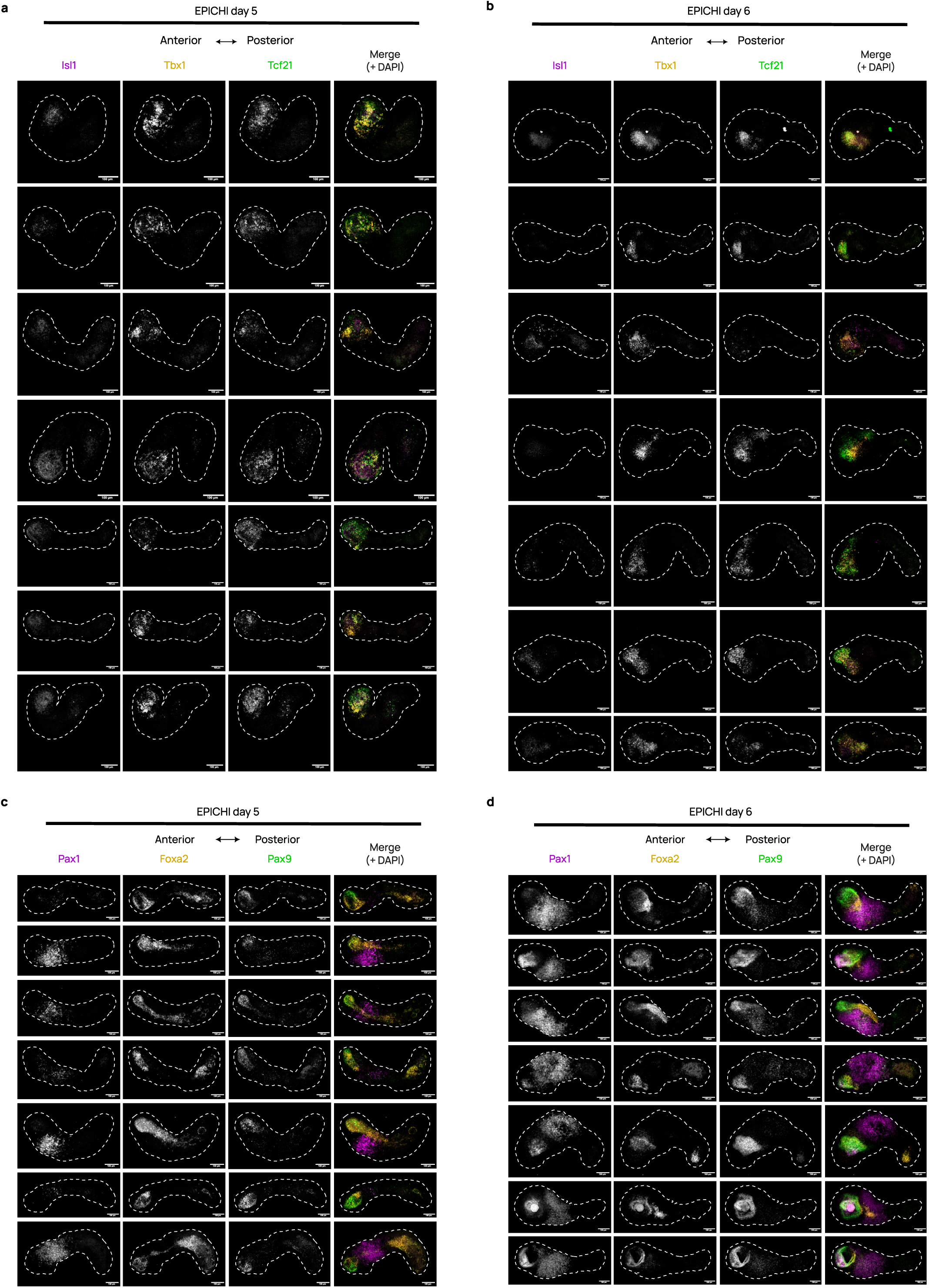
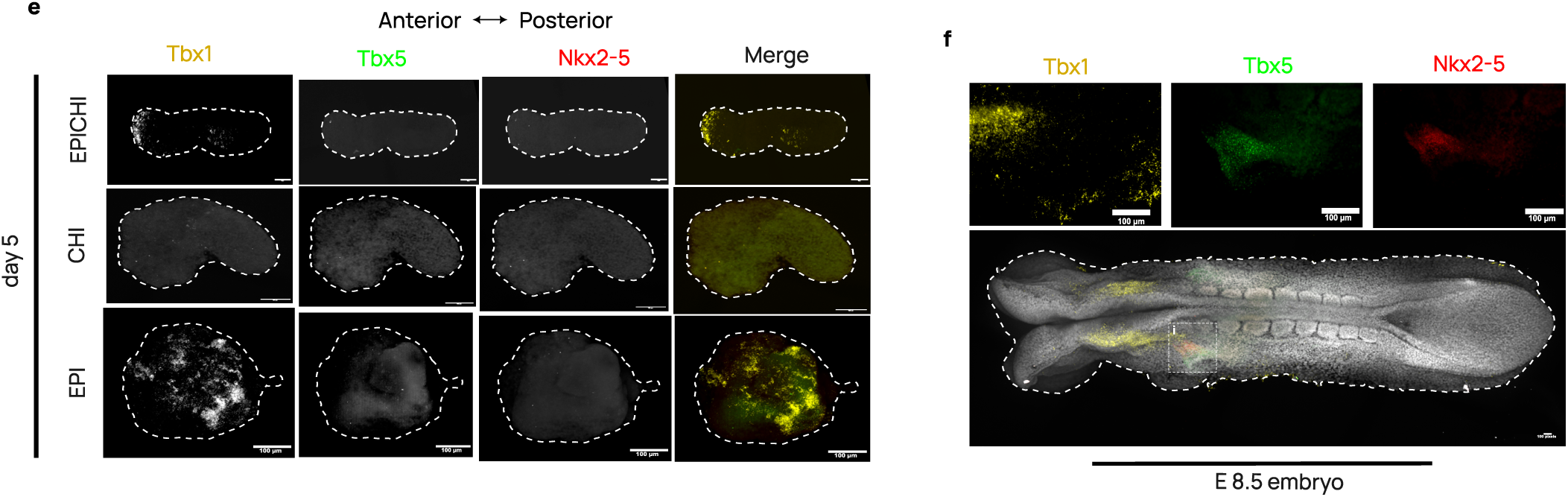
Reproducibility of pharyngeal endoderm and cardiopharyngeal mesoderm marker expression in EPICHIs. (**a, b**) EPICHIs stained with mRNA HCR probes for *Isl1* (magenta), *Tbx1* (orange), and *Tcf21* (green). Seven independent samples are shown per timepoint, at d5 (a) and d6 (b). A-P axis in left-right orientation. **(c, d)** EPICHIs stained with mRNA HCR probes for *Pax1* (magenta), *Foxa2* (orange), and *Pax9* (green). Seven independent samples are shown per timepoint, at d5 (c) and d6 (d). **(e)** Day 5 EPICHIs, CHIs and EPIs stained with mRNA HCR probes for *Tbx1* (orange), *Tbx5* (green), and *Nkx2-5* (red). *Tbx1* was detected in the anterior region of EPICHIs and broadly in EPIs, and was largely absent from CHIs; *Tbx5* and *Nkx2-5* remained undetectable in all three conditions at d5. **(f)** E8.5 mouse embryo stained with mRNA HCR probes for *Tbx1* (orange), *Tbx5* (green), and *Nkx2-5* (red). Top: detail of boxed cardiac region; bottom: whole-mount maximum intensity projection with Hoechst (grey). *Tbx5* and *Nkx2-5* mark the cardiac region adjacent to *Tbx1*⁺ pharyngeal mesoderm. Scales: 100 μm.

**Figure S8.**
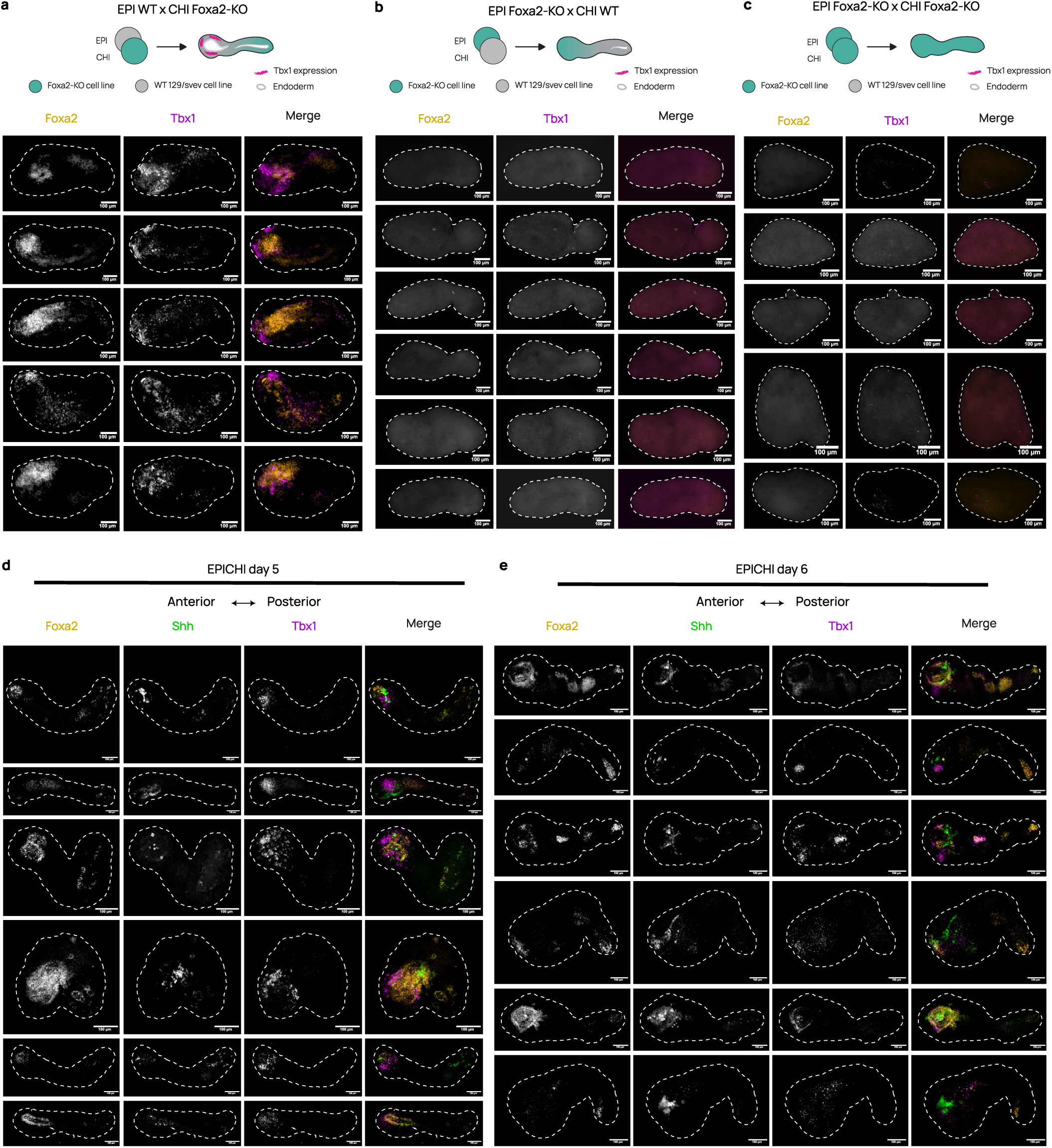
Reproducibility of *Foxa2*-dependent *Tbx1* phenotype and of *Shh* expression in EPICHI foregut epithelium. (**a-c**) Day 5 Foxa2-KO reciprocal chimeras stained with mRNA HCR probes for *Foxa2* (orange) and *Tbx1* (magenta): (a) EPI WT x CHI Foxa2-KO, (b) EPI Foxa2-KO x CHI WT, (c) EPICHI Foxa2-KO. Five to six independent samples are shown per configuration, with corresponding merging scheme above each panel. A-P axis in left-right orientation. *Foxa2* and *Tbx1* expression was retained in the anterior part when the knockout was restricted to CHI (a), and lost when the knockout affected EPI (b) or both aggregates (c). **(d, e)** EPICHIs stained with mRNA HCR probes for *Foxa2* (orange), *Shh* (green), and *Tbx1* (magenta) at d5 (d) and d6 (e). Six independent samples are shown per timepoint. *Shh* was expressed within the anterior *Foxa2*⁺ epithelium, adjacent to *Tbx1*⁺ mesoderm. Scales: 100 μm.

## REFERENCES

1. Alzamrooni, A., Mendes Vieira, P., Murciano, N., Wolton, M., Schubert, F.R., Robson, S.C., and Dietrich, S. (2023). Cardiac competence of the paraxial head mesoderm fades concomitant with a shift towards the head skeletal muscle programme. Dev. Biol. 501, 39–59. 10.1016/j.ydbio.2023.06.005.

2. Argiro, L., Chevalier, C., Choquet, C., Nandkishore, N., Ghata, A., Baudot, A., Zaffran, S., and Lescroart, F. (2024). Gastruloids are competent to specify both cardiac and skeletal muscle lineages. Nat. Commun. 15, 10172. 10.1038/s41467-024-54466-w.

3. Arias, A.M., Marikawa, Y., and Moris, N. (2022). Gastruloids: pluripotent stem cell models of mammalian gastrulation and embryo engineering. Dev. Biol. 488, 35–46. 10.1016/j.ydbio.2022.05.002.

4. Balaskas, A., Kraus, I., Özgüldez, H.Ö., Omgba, P.A., Bolondi, A., Berlad, I., Hanna, J.H., Kretzmer, H., and Bulut, A. (2025). An advanced head-to-tail mouse embryo model with hypoxia-mediated neural patterning. bioRxiv. 10.1101/2025.06.17.660116.

5. Banerjee, A., Yallapragada, S.K., Torregrosa-Cortés, G., Vyas, B.J., and Sambasivan, R. (2025). Retinoic acid coordinates the orderly construction of the mammalian body in the anterior-to-posterior sequence. 10.64898/2025.12.11.693671.

6. Beccari, L., Moris, N., Girgin, M., Turner, D.A., Baillie-Johnson, P., Cossy, A.-C., Lutolf, M.P., Duboule, D., and Arias, A.M. (2018). Multi-axial self-organization properties of mouse embryonic stem cells into gastruloids. Nature 562, 272–276. 10.1038/s41586-018-0578-0.

7. Brons, I.G.M., Smithers, L.E., Trotter, M.W.B., Rugg-Gunn, P., Sun, B., Chuva De Sousa Lopes, S.M., Howlett, S.K., Clarkson, A., Ahrlund-Richter, L., Pedersen, R.A., et al. (2007). Derivation of pluripotent epiblast stem cells from mammalian embryos. Nature 448, 191–195. 10.1038/nature05950.

8. Burtscher, I., and Lickert, H. (2009). Foxa2 regulates polarity and epithelialization in the endoderm germ layer of the mouse embryo. Development 136, 1029–1038. 10.1242/dev.028415.

9. Burtscher, I., Barkey, W., and Lickert, H. (2013). Foxa2 – venus fusion reporter mouse line allows live – cell analysis of endoderm – derived organ formation. Genesis 51, 596–604. 10.1002/dvg.22404.

10. Chang, C.-N., and Kioussi, C. (2018). Location, location, location: signals in muscle specification. J. Dev. Biol. 6, 11. 10.3390/jdb6020011.

11. Choi, H.M.T., Schwarzkopf, M., Fornace, M.E., Acharya, A., Artavanis, G., Stegmaier, J., Cunha, A., and Pierce, N.A. (2018). Third-generation *in situ* hybridization chain reaction: multiplexed, quantitative, sensitive, versatile, robust. Development 145, dev165753. 10.1242/dev.165753.

12. Costello, I., Pimeisl, I.-M., Dräger, S., Bikoff, E.K., Robertson, E.J., and Arnold, S.J. (2011). The T-box transcription factor Eomesodermin acts upstream of Mesp1 to specify cardiac mesoderm during mouse gastrulation. Nat. Cell Biol. 13, 1084–1091. 10.1038/ncb2304.

13. Dias, A., Pascual-Mas, P., Torregrosa-Cortés, G., McNamara, H.M., Wehmeyer, A.E., Arnold, S.J., and Martinez Arias, A. (2025). Opposing Nodal and Wnt signalling activities govern the emergence of the mammalian body plan. 10.1101/2025.01.11.632562.

14. Duarte, P., Brattig Correia, R., Nóvoa, A., and Mallo, M. (2023). Regulatory changes associated with the head to trunk developmental transition. BMC Biol. 21, 170. 10.1186/s12915-023-01675-2.

15. Dunn, N.R., Vincent, S.D., Oxburgh, L., Robertson, E.J., and Bikoff, E.K. (2004). Combinatorial activities of Smad2 and Smad3 regulate mesoderm formation and patterning in the mouse embryo. Development 131, 1717–1728. 10.1242/dev.01072.

16. Farag, N., Sacharen, C., Avni, L., and Nachman, I. (2024). Coordination between endoderm progression and mouse gastruloid elongation controls endodermal morphotype choice. Dev. Cell 59, 2364–2374.e4. 10.1016/j.devcel.2024.05.017.

17. Garg, V., Yamagishi, C., Hu, T., Kathiriya, I.S., Yamagishi, H., and Srivastava, D. (2001). Tbx1, a DiGeorge Syndrome Candidate Gene, Is Regulated by Sonic Hedgehog during Pharyngeal Arch Development. Dev. Biol. 235, 62–73. 10.1006/dbio.2001.0283.

18. Girgin, M.U., Broguiere, N., Mattolini, L., and Lutolf, M.P. (2021). Gastruloids generated without exogenous wnt activation develop anterior neural tissues. Stem Cell Rep. 16, 1143–1155. 10.1016/j.stemcr.2021.03.017.

19. Graham, A., and Smith, A. (2000). Patterning the pharyngeal arches. Bioessays 23, 54–61. 10.1002/1521-1878(200101)23:1%3C54::AID-BIES1007%3E3.0.CO;2-5.

20. Guzzetta, A., Koska, M., Rowton, M., Sullivan, K.R., Jacobs-Li, J., Kweon, J., Hidalgo, H., Eckart, H., Hoffmann, A.D., Back, R., et al. (2020). Hedgehog–FGF signaling axis patterns anterior mesoderm during gastrulation. Proc. Natl. Acad. Sci. 117, 15712–15723. 10.1073/pnas.1914167117.

21. Hashmi, A., Tlili, S., Perrin, P., Lowndes, M., Peradziryi, H., Brickman, J.M., Martínez Arias, A., and Lenne, P.-F. (2022). Cell-state transitions and collective cell movement generate an endoderm-like region in gastruloids. eLife 11, e59371. 10.7554/eLife.59371.

22. Hayashi, K., Ohta, H., Kurimoto, K., Aramaki, S., and Saitou, M. (2011). Reconstitution of the mouse germ cell specification pathway in culture by pluripotent stem cells. Cell 146, 519–532. 10.1016/j.cell.2011.06.052.

23. Ikonomou, L., and Kotton, D.N. (2015). Derivation of endodermal progenitors from pluripotent stem cells. J. Cell. Physiol. 230, 246–258. 10.1002/jcp.24771.

24. Kaufman-Francis, K., Goh, H.N., Kojima, Y., Studdert, J.B., Jones, V., Power, M.D., Wilkie, E., Teber, E., Loebel, D.A.F., and Tam, P.P.L. (2014). Differential response of epiblast stem cells to nodal and activin signalling: a paradigm of early endoderm development in the embryo. Philos. Trans. R. Soc. B-Biol. Sci. 369, 20130550. 10.1098/rstb.2013.0550.

25. Kinder, S.J., Tsang, T.E., Quinlan, G.A., Hadjantonakis, A.-K., Nagy, A., and Tam, P.P.L. (1999). The orderly allocation of mesodermal cells to the extraembryonic structures and the anteroposterior axis during gastrulation of the mouse embryo. Development 126, 4691–4701. 10.1242/dev.126.21.4691.

26. Kojima, Y., Kaufman-Francis, K., Studdert, J.B., Steiner, K.A., Power, M.D., Loebel, D.A.F., Jones, V., Hor, A., de Alencastro, G., Logan, G.J., et al. (2014). The transcriptional and functional properties of mouse epiblast stem cells resemble the anterior primitive streak. Cell Stem Cell 14, 107–120. 10.1016/j.stem.2013.09.014.

27. Lescroart, F., Dumas, C.E., Adachi, N., and Kelly, R.G. (2022). Emergence of heart and branchiomeric muscles in cardiopharyngeal mesoderm. Exp. Cell Res. 410, 112931. 10.1016/j.yexcr.2021.112931.

28. Lin, L., Bu, L., Cai, C.-L., Zhang, X., and Evans, S. (2006). Isl1 is upstream of sonic hedgehog in a pathway required for cardiac morphogenesis. Dev. Biol. 295, 756–763. 10.1016/j.ydbio.2006.03.053.

29. Litingtung, Y., Lei, L., Westphal, H., and Chiang, C. (1998). Sonic hedgehog is essential to foregut development. Nat. Genet. 20, 58–61. 10.1038/1717.

30. Liu, Z., Qiu, C., Kubo, C.A., Xu, S., Daza, R.M., Nichols, E., Yang, W., Vo, A., O’Neill, M.B., Lee, C., et al. (2025). Dual-patterned pluripotent stem cells self-organize into a human embryo model with extended anterior-posterior patterning. bioRxiv. 10.1101/2025.09.25.678678.

31. Martinez Arias, A., Rivron, N., Moris, N., Tam, P., Alev, C., Fu, J., Hadjantonakis, A.-K., Hanna, J.H., Minchiotti, G., Pourquie, O., et al. (2024). Criteria for the standardization of stem-cell-based embryo models. Nat. Cell Biol. 26, 1625–1628. 10.1038/s41556-024-01492-x.

32. Matthews, K.R.W., Wagner, D.S., and Warmflash, A. (2021). Stem cell-based models of embryos: The need for improved naming conventions. Stem Cell Rep. 16, 1014–1020. 10.1016/j.stemcr.2021.02.018.

33. Miao, Y., Djeffal, Y., De Simone, A., Zhu, K., Lee, J.G., Lu, Z., Silberfeld, A., Rao, J., Tarazona, O.A., Mongera, A., et al. (2023). Reconstruction and deconstruction of human somitogenesis in vitro. Nature 614, 500–508. 10.1038/s41586-022-05655-4.

34. Moncaut, N., Cross, J.W., Siligan, C., Keith, A., Taylor, K., Rigby, P.W.J., and Carvajal, J.J. (2012). Musculin and TCF21 coordinate the maintenance of myogenic regulatory factor expression levels during mouse craniofacial development. Development 139, 958–967. 10.1242/dev.068015.

35. Moris, N., Anlas, K., Van Den Brink, S.C., Alemany, A., Schröder, J., Ghimire, S., Balayo, T., Van Oudenaarden, A., and Martinez Arias, A. (2020). An in vitro model of early anteroposterior organization during human development. Nature 582, 410–415. 10.1038/s41586-020-2383-9.

36. Nandkishore, N., Vyas, B., Javali, A., Ghosh, S., and Sambasivan, R. (2018). Divergent early mesoderm specification underlies distinct head and trunk muscle programmes in vertebrates. Development 145, dev160945. 10.1242/dev.160945.

37. Parameswaran, M., and Tam, P.P.L. (1995). Regionalisation of cell fate and morphogenetic movement of the mesoderm during mouse gastrulation. Dev. Genet. 17, 16–28. 10.1002/dvg.1020170104.

38. Piotrowski, T., and Nüsslein-Volhard, C. (2000). The endoderm plays an important role in patterning the segmented pharyngeal region in zebrafish (danio rerio). Dev. Biol. 225, 339–356. 10.1006/dbio.2000.9842.

39. Probst, S., Sagar, Tosic J., Schwan, C., Grün, D., and Arnold, S.J. (2021). Spatiotemporal sequence of mesoderm and endoderm lineage segregation during mouse gastrulation. Development 148, dev193789. 10.1242/dev.193789.

40. Robertson, E.J. (2014). Dose-dependent Nodal/Smad signals pattern the early mouse embryo. Semin. Cell Dev. Biol. 32, 73–79. 10.1016/j.semcdb.2014.03.028.

41. Rossi, G., Broguiere, N., Miyamoto, M., Boni, A., Guiet, R., Girgin, M., Kelly, R.G., Kwon, C., and Lutolf, M.P. (2021). Capturing cardiogenesis in gastruloids. Cell Stem Cell 28, 230–240.e6. 10.1016/j.stem.2020.10.013.

42. Rossi, G., Giger, S., Hübscher, T., and Lutolf, M.P. (2022). Gastruloids as in vitro models of embryonic blood development with spatial and temporal resolution. Sci. Rep. 12, 13380. 10.1038/s41598-022-17265-1.

43. Sherwood, R.I., Chen, T.-Y.A., and Melton, D.A. (2009). Transcriptional dynamics of endodermal organ formation. Dev. Dyn. 238, 29–42. 10.1002/dvdy.21810.

44. Shih, H.P., Gross, M.K., and Kioussi, C. (2008). Muscle development: Forming the head and trunk muscles. Acta Histochem. 110, 97–108. 10.1016/j.acthis.2007.08.004.

45. Tam, P.P.L., Parameswaran, M., Kinder, S.J., and Weinberger, R.P. (1997). The allocation of epiblast cells to the embryonic heart and other mesodermal lineages: the role of ingression and tissue movement during gastrulation. Development 124, 1631–1642. 10.1242/dev.124.9.1631.

46. Tesar, P.J., Chenoweth, J.G., Brook, F.A., Davies, T.J., Evans, E.P., Mack, D.L., Gardner, R.L., and McKay, R.D.G. (2007). New cell lines from mouse epiblast share defining features with human embryonic stem cells. Nature 448, 196–199. 10.1038/nature05972.

47. Turner, D.A., and Nichols, J. (2023). Modifying gastruloids to dissect mechanisms of tissue-specific induction. Curr. Opin. Genet. Dev. 83, 102130. 10.1016/j.gde.2023.102130.

48. Turner, D.A., Rué, P., Mackenzie, J.P., Davies, E., and Martinez Arias, A. (2014). Brachyury cooperates with wnt/β-catenin signalling to elicit primitive-streak-like behaviour in differentiating mouse embryonic stem cells. BMC Biol. 12, 63. 10.1186/s12915-014-0063-7.

49. Tzahor, E., and Evans, S.M. (2011). Pharyngeal mesoderm development during embryogenesis: implications for both heart and head myogenesis. Cardiovasc. Res. 91, 196–202. 10.1093/cvr/cvr116.

50. Van Den Brink, S.C., and Van Oudenaarden, A. (2021). 3D gastruloids: a novel frontier in stem cell-based in vitro modeling of mammalian gastrulation. Trends Cell Biol. 31, 747–759. 10.1016/j.tcb.2021.06.007.

51. Van Den Brink, S.C., Baillie-Johnson, P., Balayo, T., Hadjantonakis, A.-K., Nowotschin, S., Turner, D.A., and Martinez Arias, A. (2014). Symmetry breaking, germ layer specification and axial organisation in aggregates of mouse embryonic stem cells. Development 141, 4231–4242. 10.1242/dev.113001.

52. Van Den Brink, S.C., Alemany, A., Van Batenburg, V., Moris, N., Blotenburg, M., Vivié, J., Baillie-Johnson, P., Nichols, J., Sonnen, K.F., Martinez Arias, A., et al. (2020). Single-cell and spatial transcriptomics reveal somitogenesis in gastruloids. Nature 582, 405–409. 10.1038/s41586-020-2024-3.

53. Veenvliet, J.V., Bolondi, A., Kretzmer, H., Haut, L., Scholze-Wittler, M., Schifferl, D., Koch, F., Guignard, L., Kumar, A.S., Pustet, M., et al. (2020). Mouse embryonic stem cells self-organize into trunk-like structures with neural tube and somites. Science 370, eaba4937. 10.1126/science.aba4937.

54. Vincent, S.D., Dunn, N.R., Hayashi, S., Norris, D.P., and Robertson, E.J. (2003). Cell fate decisions within the mouse organizer are governed by graded Nodal signals. Genes Dev. 17, 1646–1662. 10.1101/gad.1100503.

55. Wehmeyer, A.E., Schüle, K.M., Conrad, A., Schröder, C.M., Probst, S., and Arnold, S.J. (2022). Chimeric 3D gastruloids – a versatile tool for studies of mammalian peri-gastrulation development. Development 149, dev200812. 10.1242/dev.200812.

56. Wehmeyer, A.E., Schmitt, J.K., Eggersdorfer, F., Zissel, L., Schröder, C.M., Tekman, M., Dias, A., Schüle, K.M., Martinez-Arias, A., McDole, K., et al. (2025). Competing regulatory modules control the transition between mammalian gastrulation modes. bioRxiv. 10.1101/2025.05.07.652670.

57. Xu, P.-F., Borges, R.M., Fillatre, J., De Oliveira-Melo, M., Cheng, T., Thisse, B., and Thisse, C. (2021). Construction of a mammalian embryo model from stem cells organized by a morphogen signalling centre. Nat. Commun. 12, 3277. 10.1038/s41467-021-23653-4.

58. Yamagishi, H., Maeda, J., Hu, T., McAnally, J., Conway, S.J., Kume, T., Meyers, E.N., Yamagishi, C., and Srivastava, D. (2003). *Tbx1* is regulated by tissue-specific forkhead proteins through a common Sonic hedgehog-responsive enhancer. Genes Dev. 17, 269–281. 10.1101/gad.1048903.

59. Yamanaka, Y., Hamidi, S., Yoshioka-Kobayashi, K., Munira, S., Sunadome, K., Zhang, Y., Kurokawa, Y., Ericsson, R., Mieda, A., Thompson, J.L., et al. (2023). Reconstituting human somitogenesis in vitro. Nature 614, 509–520. 10.1038/s41586-022-05649-2.

